# Single-cell epigenomic identification of inherited risk loci in Alzheimer’s and Parkinson’s disease

**DOI:** 10.1101/2020.01.06.896159

**Authors:** M. Ryan Corces, Anna Shcherbina, Soumya Kundu, Michael J. Gloudemans, Laure Frésard, Jeffrey M. Granja, Bryan H. Louie, Shadi Shams, S. Tansu Bagdatli, Maxwell R. Mumbach, Bosh Liu, Kathleen S. Montine, William J. Greenleaf, Anshul Kundaje, Stephen B. Montgomery, Howard Y. Chang, Thomas J. Montine

## Abstract

Genome-wide association studies (GWAS) have identified thousands of variants associated with disease phenotypes. However, the majority of these variants do not alter coding sequences, making it difficult to assign their function. To this end, we present a multi-omic epigenetic atlas of the adult human brain through profiling of the chromatin accessibility landscapes and three-dimensional chromatin interactions of seven brain regions across a cohort of 39 cognitively healthy individuals. Single-cell chromatin accessibility profiling of 70,631 cells from six of these brain regions identifies 24 distinct cell clusters and 359,022 cell type-specific regulatory elements, capturing the regulatory diversity of the adult brain. We develop a machine learning classifier to integrate this multi-omic framework and predict dozens of functional single nucleotide polymorphisms (SNPs), nominating gene and cellular targets for previously orphaned GWAS loci. These predictions both inform well-studied disease-relevant genes, such as *BIN1* in microglia for Alzheimer’s disease (AD) and reveal novel gene-disease associations, such as *STAB1* in microglia and *MAL* in oligodendrocytes for Parkinson’s disease (PD). Moreover, we dissect the complex inverted haplotype of the *MAPT* (encoding tau) PD risk locus, identifying ectopic enhancer-gene contacts in neurons that increase *MAPT* expression and may mediate this disease association. This work greatly expands our understanding of inherited variation in AD and PD and provides a roadmap for the epigenomic dissection of noncoding regulatory variation in disease.

## INTRODUCTION

Alzheimer’s disease (AD) and Parkinson’s disease (PD) affect ∼50 and ∼10 million individuals world-wide, as two of the most common neurodegenerative disorders. Several large consortia have assembled genome-wide association studies (GWAS) that associate genetic variants with clinical diagnoses of probable AD dementia^1–4^ or probable PD^5–7^, or with their characteristic pathologic features. These efforts have led to the identification of dozens of potential risk loci for these prevalent neurodegenerative diseases. One goal of these studies was to build more precise molecular biomarkers of AD or PD, efforts that are beginning to yield encouraging results with polygenic risk scores^8^. The other major goal was to gain deeper insight into the molecular pathogenesis of disease and thereby inform novel therapeutic targets. Some of the risk loci contain coding variants and so have credibility as putative disease mediators. However, most risk loci are in noncoding regions and so it remains unclear if the nominated (often nearest) gene is the functional disease-relevant gene, or if some other gene is involved^9^. Furthermore, even if the nominated gene is a true positive, the noncoding risk locus might regulate additional genes. These challenges remain a fundamental gap in interpreting the etiology of neurodegenerative diseases and detecting high-confidence therapeutic targets.

To an extent not achieved in other organs, human brain function is closely coupled to region and thus cellular composition. However, GWAS are agnostic to the regional and cellular heterogeneity of the brain, making it difficult to *a priori* predict which brain regions or specific cell types may mediate the phenotypic association. In addition, functional noncoding SNPs would be predicted to exert their effects through alteration of gene expression via perturbation of transcription factor binding and regulatory element function^9^. Moreover, such regulatory elements are highly cell type-specific^10^. Thus, comprehensive nomination of putative functional noncoding SNPs in the brain requires cataloging the regulatory elements that are active in every brain cell type in the correct organismal and regional context. These critical data will illuminate the functional significance of genetic risk loci in the molecular pathogenesis of common neurodegenerative diseases.

Here, we have further expanded upon the current understanding of inherited variation in neurodegenerative disease through implementation of a multi-omic framework that enables accurate prediction of functional noncoding SNPs. This framework layers bulk Assay for Transposase-accessible chromatin using sequencing (ATAC-seq)^11^, single-cell ATAC-seq (scATAC-seq)^12^, and HiChIP enhancer connectome^13, 14^ data over a machine learning classifier to predict putative functional SNPs driving association with neurodegenerative diseases. Through these efforts, we pinpoint putative target genes and cell types of several noncoding GWAS locus in AD and PD, enabling the identification of putative driver polymorphisms regulating expression of key disease-relevant genes and nominating novel gene-cell type associations. Moreover, our integrative framework provides a roadmap for application of this data and technology to any neurological disorder, thus enabling a more comprehensive understanding of the role or inherited noncoding variation in disease.

## RESULTS

### Chromatin accessibility landscapes identify brain regional epigenomic heterogeneity

We profiled the chromatin accessibility landscapes of 7 brain regions across 39 cognitively healthy individuals to deeply characterize the role of the noncoding genome in neurodegenerative diseases (Supplementary Table 1). These brain regions include distinct isocortical regions [superior and middle temporal gyri (SMTG, Brodmann areas 21 and 22), parietal lobe (PARL, Brodmann area 39), and middle frontal gyrus (MDFG, Brodmann area 9)], striatum at the level of the anterior commissure [caudate nucleus (CAUD) and putamen (PTMN)], hippocampus (HIPP) at the level of the lateral geniculate nucleus, and the substantia nigra (SUNI) at the level of the red nucleus (Figure 1a). These regions were chosen to represent the diversity of brain functionality and cell type composition, and to be the most relevant to prevalent neurodegenerative diseases. In total, we generated 268 ATAC-seq libraries from 140 macrodissected brain samples, with technical replicates for 128 of the 140 samples. From these 268 ATAC-seq libraries, we compiled a merged set of 186,559 peaks reproducible across at least 30% of samples within a given brain region (Figure 1b and Supplementary Table 2; see Methods). Dimensionality reduction via t-distributed stochastic neighbor embedding (t-SNE) identified 4 distinct clusters of samples, grouped roughly by the major brain region (isocortex, striatum, hippocampus, and substantia nigra; Figure 1c). Similar groupings were observed in principal component analysis with nearly 40% of the variance explaining the difference between striatal and non-striatal brain regions (Supplementary Fig 1a-b). These samples showed no clustering based on covariates such as biological sex, post-mortem interval, or *APOE* genotype (Supplementary Fig 1c-d and Supplementary Table 1). Originally, the samples in this cohort were selected from two clinically similar but pathologically distinct research participants: (i) cognitively normal individuals with no or low neuropathological features of AD, or (ii) cognitively normal individuals with intermediate or high burden of neuropathological features of AD^15, 16^. Comparison of these clinico-pathologically normal and clinically resilient donor subgroups showed no statistically significant differences in bulk chromatin accessibility in any of the brain regions profiled (Supplementary Fig. 1e). The variability across these donor subgroups was minimal in comparison to the differences in chromatin accessibility observed across different brain regions (Supplementary Fig. 1f). For this reason, these donor subgroups were treated as a single group in the remainder of analyses.

**Figure 1.**
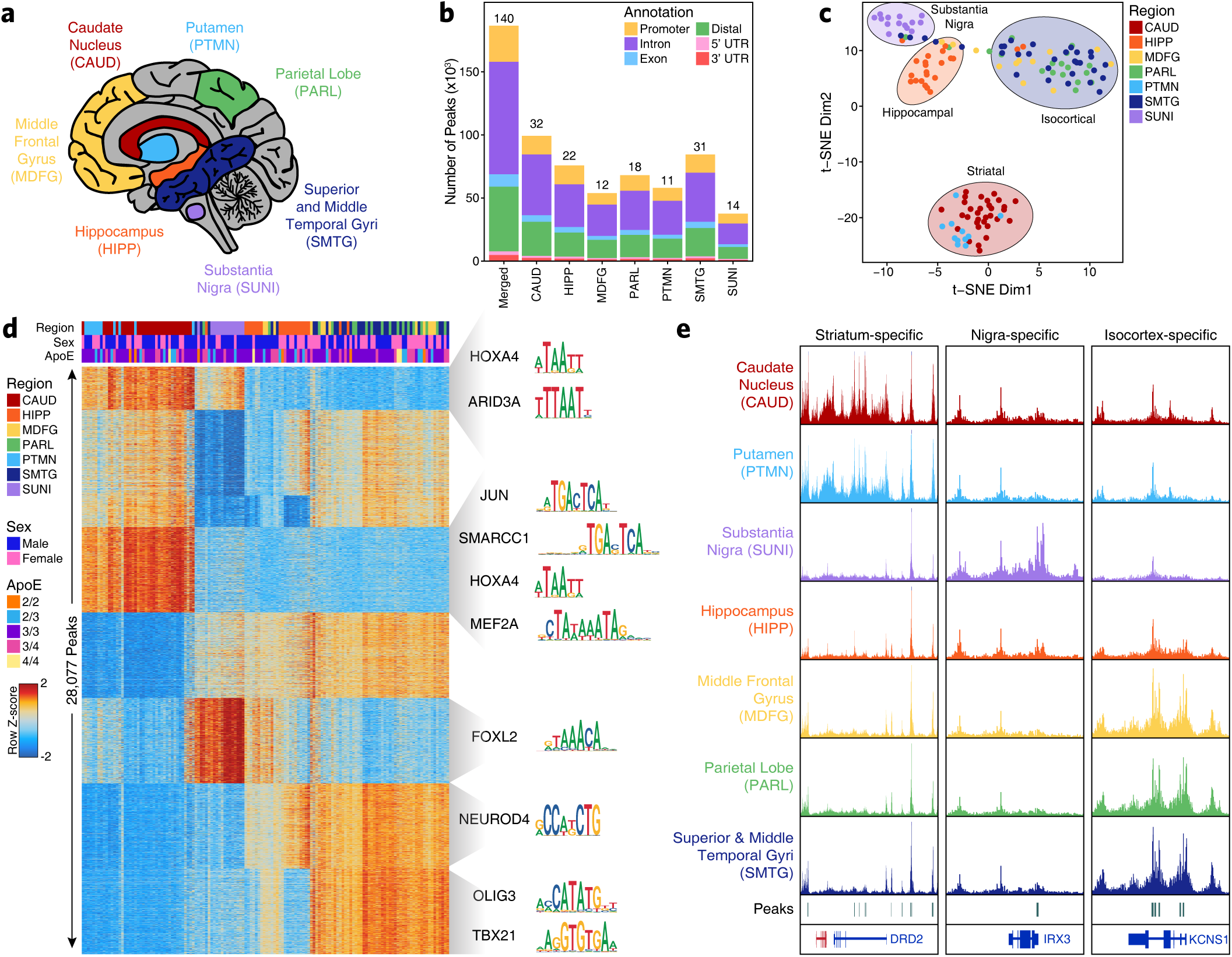
ATAC-seq defines brain-regional epigenetic heterogeneity. A. Schematic of the brain regions profiled in this study. Indicated colors are used throughout. B. Bar plot showing the number of reproducible peaks identified from samples in each brain region. The “Merged” bar represents the final merged peak set used for all bulk ATAC-seq analyses. Colors represent the type of genomic region overlapped by a given peak. The numbers above each bar represent the total number of biological samples profiled for each brain region. C. t-SNE dimensionality reduction showing all samples profiled in this study, colored by the region of the brain from which the data was generated. Each dot represents a single piece of tissue with technical replicates merged where applicable. D. Heatmap representation of binarized peaks from ATAC-seq data. Each row represents an individual peak and each column represents an individual sample. Feature groups containing more than 1000 peaks are randomly subsetted down to 1000 peaks for display on the heatmap. Feature groups containing fewer than 50 peaks are not displayed. Heatmap color represents the row-wise Z-score of normalized chromatin accessibility at the peak region. Motif names and logos shown to the right of the plot represent motifs enriched in the various peak sets. E. Sequencing tracks of region-specific ATAC-seq peaks identified through feature binarization. From left to right, *DRD2* (striatum-specific; chr11:113367951-113538919), *IRX3* (substantia nigra-specific; chr16:54276577-54291319), and *KCNS1* (isocortex-specific; chr20:45086706-45107665). Track heights are the same in each vertical panel.

Assessment of regional variation in chromatin accessibility through “feature binarization” (see Methods) identified 28,077 peaks showing region-specific or multi-region-specific accessibility (Figure 1d). For example, 14,628 and 1,734 peaks were identified with significantly increased chromatin accessibility only in striatum or substantia nigra, respectively (Figure 1d). These peak sets showed enrichment for key brain-related transcription factors (TFs) in the FOX, NEUROD, and OLIG families, consistent with suspected brain-relevant enhancers and promoters (Figure 1d). Moreover, some peaks within these sets were in the vicinity of key cell lineage-defining genes such as the dopamine receptor D2 (*DRD2*) in striatal regions, iroquois homeobox 3 (*IRX3*) in the substantia nigra, and potassium voltage-gated channel modifier subfamily S member 1 (*KCNS1*) in the isocortical regions (Figure 1e). Notably, while the hippocampus shares many peaks with other regions, we identified only 29 peaks that showed significantly increased chromatin accessibility specifically in this region. Taken together, these results indicate an extensive degree of brain regional heterogeneity that is likely representative of the functional and cellular diversity of the brain regions studied here.

### ATAC-seq refines interpretation of inherited risk variants in neurodegeneration

Using this atlas of regional chromatin accessibility, we sought to identify functional noncoding regulatory elements that may be impacted by disease-associated genetic variation identified through genome-wide association studies. Approximately 90% of phenotype-associated GWAS polymorphisms reside in noncoding DNA^17^, making it difficult to predict a putative functional impact. Moreover, linkage disequilibrium (LD) makes it difficult to pinpoint a single causative SNP when many other nearby SNPs are co-inherited. To resolve these complexities, we used a multi-tiered approach to predict which GWAS SNPs may be functional. First, we identified a compendium of SNPs that could be associated with either AD or PD (Supplementary Table 3, see Methods). To do this, we identified (i) any SNPs passing genome-wide significance in recent GWAS^1–3, 5–7^, (ii) any SNPs exhibiting colocalization of GWAS and eQTL signal, and (iii) any SNPs in linkage disequilibrium with a SNP in the previous two categories. In total, this identified 9,741 SNPs including 3,245 unique SNPs across 44 loci associated with AD and 6,496 unique SNPs across 86 loci associated with PD, with a single locus containing 34 SNPs appearing in both diseases. We then performed LD score regression to identify brain regional enrichment of neurodegeneration-related SNPs in noncoding regulatory regions. However, these regional analyses showed minimal enrichment of GWAS SNPs in peak regions associated with any of the brain regions profiled (Supplementary Fig. 2a-b). These results provide evidence against a possible regional effect involving most cell types in a particular area of the brain, but leave open the possibility of involvement of specific cell types in specific regions of the brain. Thus, we hypothesized that a single-cell-based approach could provide more granularity in identifying the precise cell types mediating disease-relevant genetic associations.

### Single-cell ATAC-seq captures regional and cell type-specific heterogeneity

To test this hypothesis and to better understand brain-regional cell type-specific chromatin accessibility landscapes, we performed single-cell chromatin accessibility profiling in 10 samples spanning the isocortex (N=3), striatum (N=3), hippocampus (N=2), and substantia nigra (N=2) (Supplementary Table 1). In total, we profiled chromatin accessibility in 70,631 individual cells (Figure 2a) after stringent quality control filtration (Supplementary Fig. 2c and Supplementary Table 4). Unbiased iterative clustering^12, 18^ of these single cells identified 24 distinct clusters (Figure 2a) which were assigned to known brain cell types based on gene activity scores (see Methods) compiled from chromatin accessibility signal in the vicinity of key lineage-defining genes^18, 19^ (Figure 2b and Supplementary Fig. 2c). For example, chromatin accessibility at the myelin associated glycoprotein (*MAG*) gene locus defined clusters corresponding to oligodendrocytes while genes such as vesicular glutamate transporter 1 (*VGLUT1* / *SLC17A7*) and vesicular GABA transporter (*VGAT* / *SLC32A1*) defined excitatory and inhibitory neurons, respectively (Figure 2b). Additionally, 13 of the 24 clusters showed regional specificity with some clusters being made up almost entirely from a single brain region (Figure 2c and Supplementary Table 4). This is most obvious for neuron, astrocyte, and oligodendrocyte precursor cell (OPC) clusters which show clear region-specific differences in clustering (Supplementary Fig. 3a-b). From this cluster-based perspective, we did not identify any clusters that were clearly segregated by gender but the sample size used in this study was not powered to make such a determination (Supplementary Fig. 3c). Cumulatively, we defined 8 distinct cell groupings and identified one cluster (Cluster 18) as putative doublets that we excluded from downstream analyses (Figure 2a and Supplementary Fig. 3d). These cell groupings varied largely in the total number of cells per grouping (Supplementary Fig. 3e) and showed distinct donor and regional compositions (Supplementary Fig. 3f-i).

**Figure 2.**
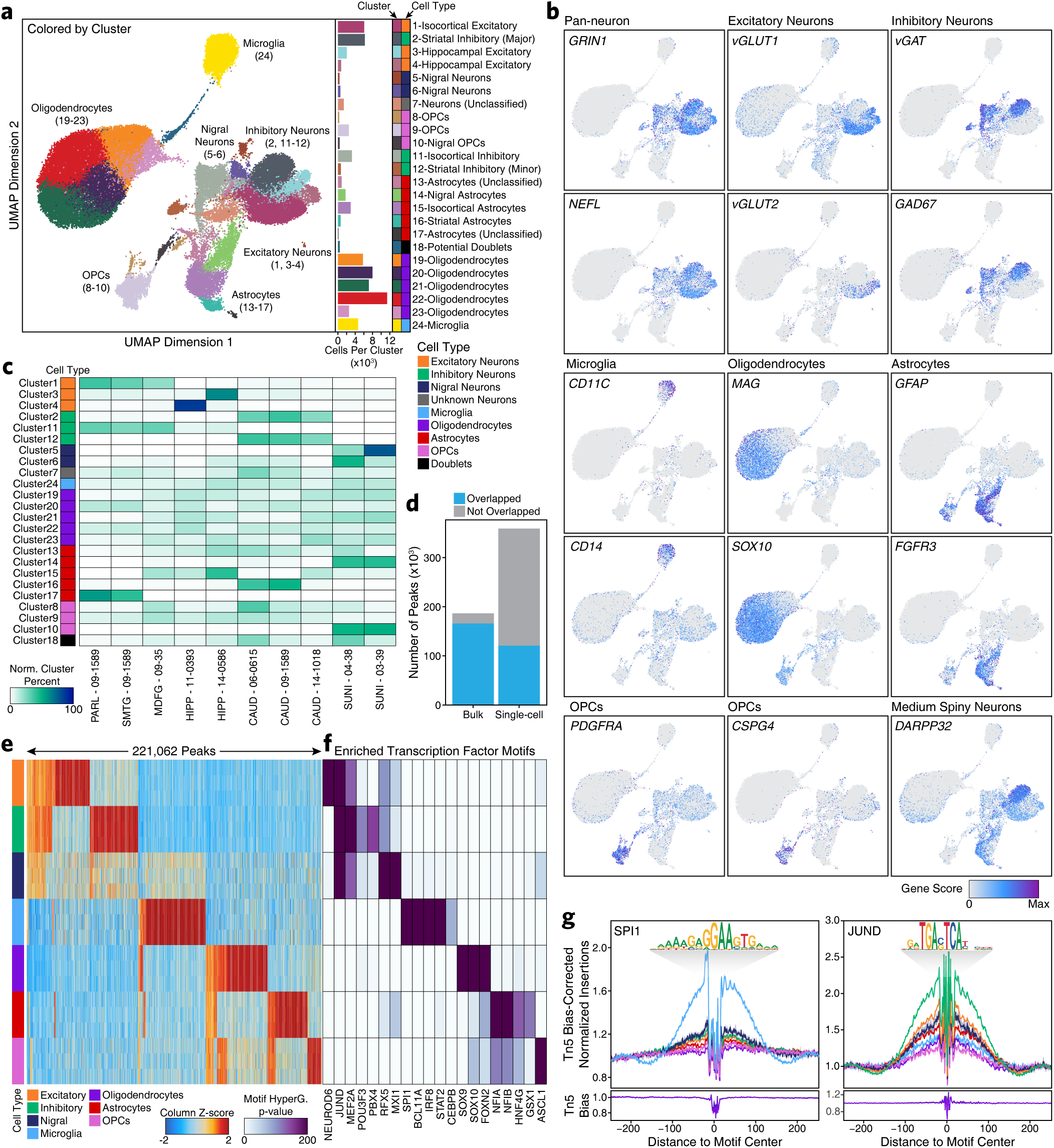
Single-cell ATAC-seq identifies cell type-specific chromatin accessibility in the adult brain. A. Left; UMAP dimensionality reduction showing identified clusters of cells. Each dot represents a single cell (N = 70,631). Right; Bar plot showing the number of cells per cluster. Each cluster is labeled to the right of the bar plot and the predicted cell type corresponding to each cluster is shown colorimetrically. B. The same UMAP dimensionality reduction shown in Figure 2a but each cell is colored by its gene activity score for the annotated lineage-defining gene. Grey represents a gene activity score of 0 while purple represents the maximum gene activity score for the given gene. C. Cluster residence heatmap showing the percent of each cluster that is composed of cells from each sample. Cell numbers were normalized across samples prior to calculating cluster residence percentages. D. Bar plot showing the overlap of bulk ATAC-seq and scATAC-seq peak calls. “Bulk” represents the number of peaks from the bulk ATAC-seq merged peak set that are overlapped by a peak called in our scATAC-seq merged peak set. “Single-cell” represents the number of peaks from our scATAC-seq merged peak set that are overlapped by a peak called in our bulk ATAC-seq merged peak set. E. Heatmap representation of binarized peaks from scATAC-seq data. Each row represents an individual pseudo-bulk replicate (3 per cell type) and each column represents an individual peak. Feature groups containing fewer than 1000 peaks are not displayed. Heatmap color represents the column-wise Z-score of normalized chromatin accessibility at the peak region. F. Motif enrichments of binarized peaks identified in Figure 2e. Due to redundancy in motifs, TF drivers were predicted using average gene expression in GTEx brain samples and accessibility at TF promoters in cell type-grouped scATAC-seq profiles. The final list of TFs represents a trimmed set of all TFs with the most likely driving TF labeled below. Color represents the p-value of the hypergeometric test for motif enrichment. G. Footprinting analysis of the SPI1 (left) and JUND (right) transcription factors across the 7 major cell types. The motif logos are shown above and the Tn5 transposase insertion biases are shown below.

Using these robustly defined clusters, we then called peaks of pseudo-bulk chromatin accessibility to create a union set of 359,022 reproducible peaks (Supplementary Table 5). Overall, 89% of the bulk ATAC-seq peaks were overlapped by a peak called in the scATAC-seq data (Figure 2d). Conversely, only 34% of the scATAC-seq peaks were overlapped by a peak from the bulk ATAC-seq peak set (Figure 2d). This is consistent with the known difficulty in identifying peaks in bulk data derived from cell types that comprise less than 20% of the total cells in the tissue^20^. These results highlight the utility of single-cell methods in situations where cell type-specific peaks are difficult to identify from bulk tissues containing multiple distinct cell types at varying frequencies.

This single-cell ATAC-seq-derived peak set enabled the identification of 221,062 highly cell type-specific peaks (Figure 2e). These peaks, comprising more than 60% of all peaks identified in our single-cell data, were selected to be specific to a single cell type or specifically shared across up to three cell types using “feature binarization” (see Methods). For example, some peaks are shared across the 3 different neuronal groups (excitatory, inhibitory, nigral) while others are shared across astrocytes, OPCs, and oligodendrocytes (Figure 2e, Supplementary Table 6). However, the majority of cell type-specific peaks are uniquely accessible in a single cell type; for example, microglia show 45,196 peaks that are specifically accessible in microglia and not in any of the other cell types profiled (Figure 2e). In total, more than 47% of the peaks called in our single-cell ATAC-seq data are specific to a single cell type (Supplementary Table 6) with the vast majority of these cell type-specific peaks remaining undetected in our bulk ATAC-seq analyses. To predict which TFs may be responsible for establishing and maintaining these cell type-specific regulatory programs, we performed motif enrichment analyses of peaks specific to each cell type (Figure 2f). We identified many known drivers of cell type identity, such as motifs specific to SOX9 and SOX10 in oligodendrocytes^21, 22^, or to ASCL1 in OPCs^23, 24^. Lastly, TF footprinting from our scATAC-seq-derived cell type-specific chromatin accessibility data showed enrichment of binding of key lineage defining TFs SPI1 and JUND in microglia and neurons, respectively (Figure 2g). Overall, these results provide a reference map of chromatin accessibility in the adult brain at single-cell resolution.

### Single-cell ATAC-seq provides reference cell populations for deconvolution of cell type-specific signals in bulk data

Using the cell type-specific signals present in our scATAC-seq data (Supplementary Fig. 4a), we performed cell type deconvolution of our bulk ATAC-seq data using CIBERSORT^25^ (Supplementary Table 7). Using our 8 cell type classification, we deconvolved the ATAC-seq signal from all 140 samples profiled by bulk ATAC-seq in this study, finding clear and expected patterns of cell type abundance such as a relative absence of excitatory neurons in the striatum (Supplementary Fig. 4b). Similarly, deconvolution based on clusters shows expected patterns including the mapping of signal from Cluster 14 (nigral astrocytes) specifically to samples from the substantia nigra, and mapping of signal from Cluster 2 (striatal inhibitory neurons) specifically to samples from the striatum (Supplementary Fig. 4c). By comparing the CIBERSORT prediction to the observed “ground truth” in the scATAC-seq data for the 10 samples profiled here, we were able to assess the performance of the cell type-specific and cluster-specific classifiers (Supplementary Fig. 4d-e). As would be expected, the cell type-specific classifier showed better performance than the cluster-specific classifier, largely due to over-or under-prediction of closely related clusters, such as the oligodendrocytic Clusters 19-23, by the cluster-specific classifier (Supplementary Fig. 4e). Application of the cell type-specific and cluster-specific classifiers to each individual bulk ATAC-seq sample profiled above showed a striking degree of variability in the bulk data based on predicted cell type abundance (Supplementary Fig. 4f-g). Such large differences in cell type composition can hamper efforts to find differential features, further supporting the use of single-cell approaches to understand complex tissues and disease states where small disease-specific variation may be overshadowed by larger differences in cell type composition across samples.

### Single-cell ATAC-seq identifies brain region-specific differences in glial cells

Our dissection of the cell type-specific chromatin landscapes in adult brain identified clusters that are both region-and cell type-specific such as Cluster 14 which is comprised almost exclusively of astrocytes from the substantia nigra (Figure 2c and Supplementary Table 4). This observation indicates that certain brain cell types may show region-specific variation. This phenomenon has been very well described in neurons, with, for example, inhibitory neurons from the striatum (largely medium spiny neurons) differing substantially from inhibitory neurons outside of the striatum^26^. Murine oligodendrocytes^27^ and astrocytes^28^ also show regional differences in morphology, function, and gene expression. However, the brain-regional variation of glial cells in humans remains less well understood. To address this, we grouped cells into one of the 8 broad cell types defined above and created pseudo-bulk reference populations from the cumulative data (see Methods). Using these region-cell type combinations, we calculated Pearson correlations for all regions across a single cell type (Supplementary Fig. 5a). As expected, neuronal cell types showed the most regional variation.

Glial cells, however, also showed substantial regional variation, with astrocytes showing the most variation followed by OPCs (Supplementary Fig. 5a). Within astrocytes, the greatest difference was found between the substantia nigra and the isocortex, indicating that the function or composition of astrocytes may differ across these brain regions. Differential peak analysis identified significant differences in chromatin accessibility near transcriptional regulators that may help explain the observed regional astrocytic differences (Supplementary Fig. 5b and Supplementary Table 8). In particular, nigral astrocytes showed significantly increased accessibility at the forkhead box B1 (*FOXB1*), *IRX1*, *IRX2*, *IRX3*, and *IRX5* genes. Conversely, isocortical astrocytes showed significantly increased accessibility at the *FOXG1*, zic family member 2 (*ZIC2*), and *ZIC5* genes. These changes in chromatin accessibility would be expected to correlate with similar changes in gene expression for the annotated genes. Moreover, the gene activity scores of these genes are definitional for the region-cell subtypes with, for example, *FOXB1* being active only in nigral astrocytes and *ZIC2* and *ZIC5* being active in all other astrocytes (Supplementary Fig. 5c-d). Of particular interest, the observed FOX switch from *FOXG1* in isocortical (and hippocampal/striatal) astrocytes to *FOXB1* in nigral astrocytes and the significant changes in chromatin accessibility at the IRX genes represent a potential transcriptional lineage control mechanism that could help to better understand region-specific functional differences in these astrocytes. Notably, diencephalic brain regions such as the substantia nigra have previously been shown to express *FOXB1*^29^, *IRX1*^30^, and *IRX3*^31^ during early brain development, thus explaining part of this broad TF-based lineage control. These transcriptional regulators could be exploited to drive differentiation programs to, for example, create regionally biased glial cells in vitro.

In addition to controlling regional astrocytic identity, chromatin accessibility at *IRX* genes was also found to differentiate nigral OPCs from isocortical OPCs (Supplementary Fig. 5d-e). Similarly, *FOXG1* also showed significantly more accessibility in isocortical OPCs, echoing the observations from astrocytes. Lastly, chromatin accessibility at the *PAX3* gene locus was significantly higher in nigral OPCs compared to isocortical OPCs (Supplementary Fig. 5d-e). Taken together, these results identify shared and disparate transcriptional regulatory programs that likely control regional differences amongst astrocytes and OPCs in the substantia nigra and isocortex.

Compared to astrocytes, oligodendrocytes and microglia showed less regional variation in chromatin accessibility (Supplementary Fig. 5f-g). While a small number of genes showed highly significant regional differences in oligodendrocytes (Supplementary Fig. 5h), very few genes showed appreciable regional differences among microglia. As noted previously, the regional differences observed in glial cells are a small fraction of the size and magnitude of regional differences observed in neurons (Supplementary Fig. 5i-j), further emphasizing the importance of single-cell approaches to study complex tissues.

### Single-cell ATAC-seq pinpoints the cellular targets of GWAS polymorphisms

Having generated high-quality cell type-specific chromatin accessibility profiles using scATAC-seq, we sought to refine our previous interpretation of GWAS polymorphisms. More specifically, we aimed to use these data to predict which cell type(s) may be the functional targets of various polymorphisms. When using peaks called in bulk ATAC-seq, we found that 78 LD-expanded SNPs in AD and 186 LD-expanded SNPs in PD overlapped peak regions. Combining our bulk ATAC-seq and scATAC-seq peak sets, we found that 438 SNPs in AD and 880 SNPs in PD directly overlapped peak regions. This represents a 5-fold increase in the number of SNPs observed to overlap peaks called from bulk ATAC-seq alone (Supplementary Table 3), illustrating the importance of cell type-specific interrogation of noncoding regions to dissect GWAS polymorphisms. Cell type-specific LD score regression using AD and PD GWAS results revealed a significant increase in per-SNP heritability for AD in the microglia peak set, reinforcing previous studies^2, 32, 33^ (Figure 3a and Supplementary Table 9). Similar analyses in PD showed no significant enrichment in SNP heritability in any particular cell type, perhaps indicating that the cellular bases of PD are more heterogeneous than AD (Figure 3a). Though not a focus of the current study, we note that the data generated here can be used to inform the cellular ontogeny of any brain-related GWAS. For example, we observe a striking enrichment of SNP heritability for schizophrenia, neuroticism, and attention deficit hyperactivity disorder in excitatory and inhibitory neurons (Figure 3a). We also confirmed that the heritability of GWAS SNPs from traits not directly related to brain cell types, such as lean body mass, were not enriched in any of the tested brain cell types and that cell types not expected to be involved in brain-related diseases show no enrichment of SNP heritability for brain-related disease SNPs (Supplementary Fig. 6a). Thus, combination of our scATAC-seq data with our curated list of disease-relevant SNPs enables prediction of the cellular targets of each polymorphism.

**Figure 3.**
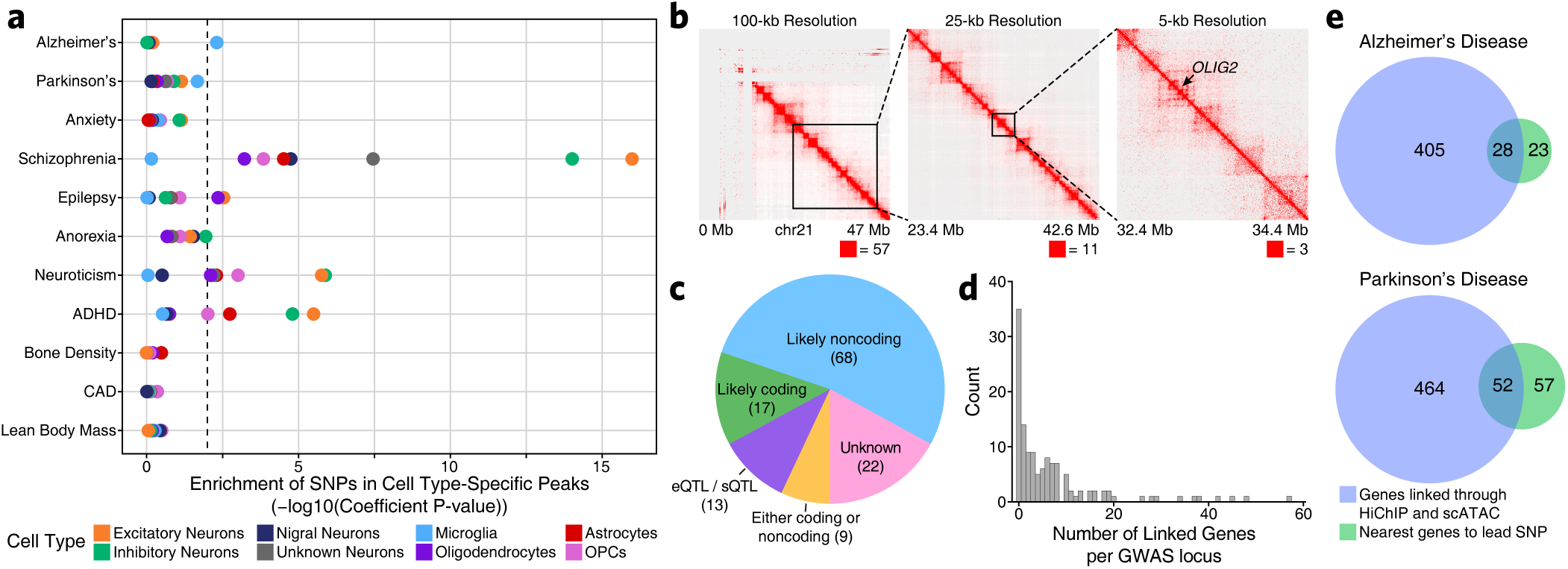
HiChIP and scATAC-seq predict gene and cellular targets of disease-associated polymorphisms. A. LD score regression identifying the enrichment of GWAS SNPs from various brain- and non-brain-related conditions in the peak regions of various cell types derived from pseudo-bulk-based scATAC-seq data. B. Heatmap representation of HiChIP interaction signal at 100-kb, 25-kb, and 5-kb resolution at the *OLIG2* locus. C. Characterization of GWAS loci in AD and PD according to the predicted effects of the polymorphisms. For example, loci whose phenotypic association is likely mediated by changes in coding regions are marked as “Likely coding”. Loci whose effect could be mediated by either coding or noncoding mechanisms are marked as “Either coding or noncoding” whereas loci with no polymorphisms overlapping a peak region or an exonic region are marked as “Unknown”. D. Histogram of the number of genes linked per GWAS locus. Each bar represents a bin of length 1. E. Venn diagram of (i) the number of genes linked through assessment of the nearest gene to the lead SNP of each AD (top) and PD (bottom) GWAS locus and (ii) the number of genes linked though HiChIP and scATAC-seq analyses of LD-expanded polymorphisms.

### Three-dimensional chromatin landscapes nominate novel target genes of inherited risk variants

In addition to understanding the cell type-specific impacts of an individual polymorphism, we also wanted to predict the gene(s) that may be the direct regulatory targets of a given noncoding polymorphism. We reasoned that the vast majority of functional GWAS SNPs would reside in noncoding sequences and therefore exert their effects through modulation of enhancer or promoter activity. As such, we mapped the enhancer-centric three-dimensional (3D) chromatin architecture in multiple brain regions using HiChIP for histone H3 lysine 27 acetylation (H3K27ac) which marks active enhancers and promoters (Figure 3b and Supplementary Fig. 6b). In total, we generated 3D interaction maps for 6 of the 7 regions profiled by ATAC-seq (putamen was excluded given the high overlap with the caudate nucleus) with an average of 158 million valid interaction pairs identified per region (Supplementary Fig. 6c). These maps led to the identification of 833,975 predicted 3D interactions across all brain regions profiled of which 331,730 (40%) were reproducible in at least two brain regions (Supplementary Fig. 6d and Supplementary Table 10). Of these loops, 29.2% had an ATAC-seq peak present in one anchor, 67.4% had an ATAC-seq peak present in both anchors, and 3.4% did not overlap any ATAC-seq peaks identified in either the bulk or scATAC-seq datasets (Supplementary Fig. 6e). Additionally, correlated variation of chromatin accessibility in peaks across single cells has been shown to predict functional interactions between regulatory elements^19, 34^. Using this co-accessibility framework, we predicted regulatory interactions from our scATAC-seq data (Supplementary Fig. 6f), identifying 2,822,924 putative interactions between regions of chromatin accessibility (Supplementary Table 10). This set of interactions showed only moderate overlap (∼20%) with our HiChIP data, consistent with the ability of this technique to identify cell type-specific regulatory interactions, whereas HiChIP of bulk brain tissue is better suited for identification of more shared regulatory interactions (Supplementary Fig. 6f). Together, these two techniques define a compendium of putative regulatory interactions in the various brain regions studied here.

To predict which genes may be altered by noncoding GWAS polymorphisms, we first classified GWAS loci according to whether their phenotypic association was likely mediated by alterations in the coding or noncoding genome (Figure 3c). Across AD and PD, this identified 17 loci that harbored likely functional coding alterations, 68 loci that harbored likely functional noncoding alterations, 9 loci that could be associated with putatively functional coding and noncoding alterations, and 22 loci that did not harbor any SNPs in coding regions nor any SNPs in regulatory regions identified in our chromatin accessibility data (Supplementary Table 3). These “unknown” loci likely represent noncoding associations in cell types that were not adequately represented in our analysis. From the original set of 9,741 disease-related SNPs, we identified 438 SNPs for AD and 880 SNPs for PD that overlapped peak regions of chromatin accessibility. Of these SNPs, 395 and 531 were involved in a putative enhancer-promoter interaction identified in our HiChIP or co-accessibility data for AD and PD, respectively (Supplementary Table 3). Cumulatively, this enabled the identification of 433 and 516 genes putatively affected by the activity of GWAS polymorphisms in AD and PD, respectively (Figure 3d-e). These gene sets are enriched for biological processes known to be implicated in AD and PD including lipoprotein particle clearance^1^ (AD) and synaptic vesicle recycling^35^ (PD) (Supplementary Fig. 6g-h).

### Machine learning predicts putative functional SNPs and identifies the molecular ontogeny of disease associations

To disentangle further the molecular underpinnings of AD and PD associations, we developed a multi-omic approach to predict functional noncoding GWAS polymorphisms (Figure 4a and Supplementary Fig. 7a). This approach is anchored in the use of a machine learning framework to score the allelic effect of a SNP on chromatin accessibility. Using the gapped *k*-mer support vector machine (gkm-SVM) framework^36^, we trained models to learn the patterns and grammars of chromatin accessibility using our scATAC-seq data (Figure 4b). Specifically, for each cluster (cell type) identified from the scATAC-seq data, we provided 1000-bp sequences centered at all of the peak regions from the cluster-specific pseudo-bulk ATAC-seq data and an equal number of GC-matched non-accessible genomic sequences to a gkm-SVM classifier and trained it to predict whether each sequence is accessible or not. The gkm-SVM models for all 24 scATAC-seq clusters exhibited high prediction performance on held-out test sequences (Supplementary Fig. 7b-c), across all folds of a 10-fold validation training paradigm (Supplementary Fig. 7d).

**Figure 4.**
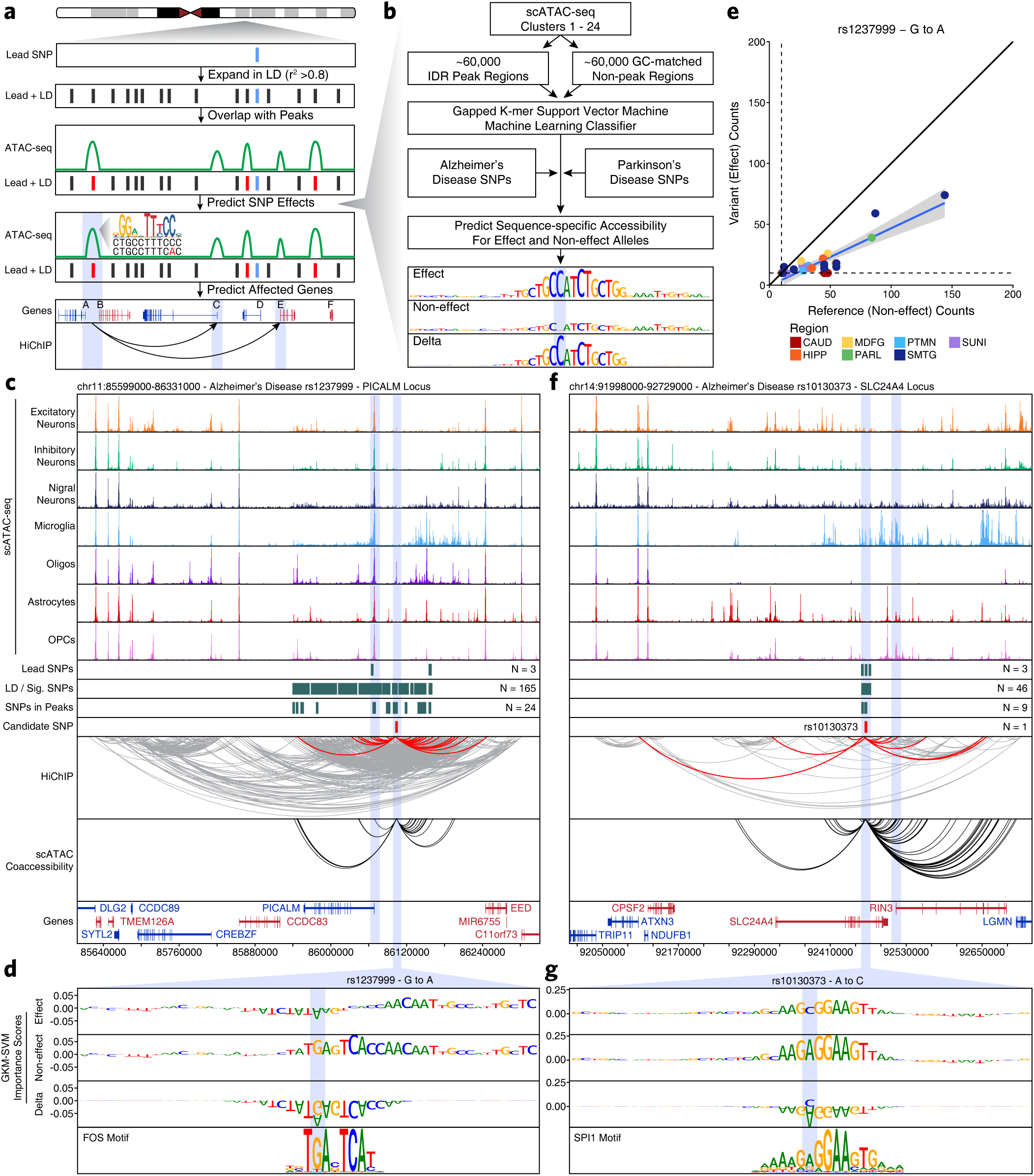
Machine learning predicts functional polymorphisms in AD and PD. A. Schematic of the overall strategy for identification of putative functional SNPs and their corresponding gene targets. B. Schematic of the gkm-SVM machine learning approach used to predict which noncoding SNPs alter transcription factor binding and chromatin accessibility. C. Normalized scATAC-seq-derived pseudo-bulk tracks, HiChIP loop calls, co-accessibility correlations, and machine learning predictions for LD-expanded SNPs in the *PICALM* gene locus. For HiChIP, each line represents a loop connecting the points on each end. Red lines contain one anchor overlapping the SNP of interest while grey lines do not. D. GkmExplain importance scores for each base in the 50-bp region surrounding rs1237999 for the effect and non-effect alleles from the gkm-SVM model corresponding to oligodendrocytes (Cluster 21). The predicted motif affected by the SNP is shown at the bottom and the SNP of interest is highlighted in blue. E. Dot plot showing allelic imbalance at rs1237999. The ATAC-seq counts for the reference/non-effect (G) allele and variant/effect (A) allele are plotted. Each dot represents an individual bulk ATAC-seq sample colored by the brain region from which the sample was collected. F. Sequencing tracks as shown in Figure 4c but for the *SLC24A4* locus. G. GkmExplain importance scores for each base in the 50-bp region surrounding rs10130373 for the effect and non-effect alleles from the gkm-SVM model corresponding to microglia (Cluster 24). The predicted motif affected by the SNP is shown at the bottom and the SNP of interest is highlighted in blue.

Next, we used three complementary approaches, GkmExplain^37^, *in silico* mutagenesis^38^, and deltaSVM^39^ to predict the allelic impact of 1677 candidate SNPs on chromatin accessibility in each cluster by providing the sequences corresponding to both alleles of each SN to the models for each of the 24 clusters. All three approaches showed high concordance of predicted allelic effects across all candidate SNPs (Supplementary Fig. 7e). In total, among the 1677 SNPs that we scored, we identified 44 high-confidence, and 41 moderate-confidence SNPs that the model predicts will have a functional consequence on chromatin accessibility via identifiable TF binding sites. Integration of these predictions with our colocalization, HiChIP, and scATAC-seq data sets allowed for a comprehensive interrogation of the epigenetic effects of noncoding polymorphisms in AD and PD (Figure 4a and Supplementary Table 3).

This multi-omic approach identifies two main categories of novel associations: established disease-related genes where the precise causative SNP remains unknown, and novel genes previously not implicated in disease pathogenesis. In each of these categories, our integrative analysis implicates SNP-gene associations that are supported by (i) the presence of the SNP in an ATAC-seq peak (Tier 3), (ii) a colocalization, HiChIP interaction, or co-accessibility correlation linking the SNP to one or more genes (Tier 2), and in many cases (iii) orthogonal prediction of SNP function via either allelic imbalance (Supplementary Fig.7f), machine learning predictions, or both (Tier 1) (Supplementary Fig. 7a). Allelic imbalance refers to the differential accessibility between two alleles when one allele is more readily bound than the other. This is obtained from our bulk ATAC-seq data which is available for all donors, thus highlighting the utility of a combined bulk and single-cell approach. Moreover, the cell type-specificity of our scATAC-seq data allows identification of the cell types in which these disease associations likely form.

Many studies have investigated the role of genes such as Phosphatidylinositol Binding Clathrin Assembly Protein (*PICALM*)^40^, Solute Carrier Family 24 Member 4 (*SLC24A4*)^41^, Bridging Integrator 1 (*BIN1*)^10, 42^, and Membrane Spanning 4-Domains A6A (*MS4A6A*)^43^ in AD since their implication in the disease by GWAS. However, it remains unclear which polymorphisms drive these associations. In the case of *PICALM*, our models predict a potential functional variant (rs1237999) which resides within an oligodendrocyte-specific regulatory element 35-kb upstream of *PICALM* and disrupts a putative FOS/AP1 factor binding site (Figure 4c-d). Moreover, rs1237999 shows striking allelic imbalance with the variant (effect) allele showing diminished accessibility in bulk ATAC-seq data from heterozygotes across multiple brain regions (Figure 4e). Lastly, rs1237999 shows 3D interaction with both *PICALM* and the *EED* gene, a polycomb-group family member involved in maintaining a repressive transcriptional state. This expands the potential functional role of this association to a novel gene and specifically points to a role for oligodendrocytes which were not previously implicated in this phenotypic association^40^.

Similarly, the *SLC24A4* locus harbors a small LD block with 46 SNPs that all reside within an intron of *SLC24A4*. Previous work has implicated both *SLC24A4* and the nearby Ras And Rab Interactor 3 (*RIN3*) gene in this association but the true mediator remains unclear^44, 45^. Our multi-omic approach identifies a single SNP, rs10130373, which occurs within a microglia-specific peak, disrupts an SPI1 motif, and communicates specifically with the promoter of the *RIN3* gene (Figure 4f-g). This is consistent with the role of *RIN3* in the early endocytic pathway which is crucial for microglial function and of particular disease relevance in AD^46^.

In the case of *BIN1*, our work and previous work^10^ predict SNP rs6733839 to disrupt a MEF2 binding site in a microglia-specific enhancer located 28-kb upstream of the *BIN1* promoter (Supplementary Fig. 8a). Our machine learning framework additionally implicates SNP rs13025717 which we predict to disrupt a KLF4 binding motif in a microglia-specific putative enhancer 21-kb upstream of *BIN1* (Supplementary Fig. 8b). Both of these SNPs have previously been shown to have sequence-specific correlations with *BIN1* gene expression^47^. Similarly, we identified rs636317 in the *MS4A6A* locus which disrupts a microglia-specific CTCF binding motif (Supplementary Fig. 8c-d). Cumulatively, these results annotate the most likely functional SNPs mediating known disease associations in AD and PD (Supplementary Table 3). Importantly, these predicted functional SNPs do not always affect the expected cell type nor target the closest gene, further emphasizing the utility of our integrative multi-omic approach.

Nevertheless, the true promise in studying these noncoding polymorphisms is the identification of novel genes affected by disease-associated variation. This is perhaps most important in PD where identification of disease-associated genes is less mature. The *ITIH1* GWAS locus occurs within a 600-kb LD block harboring 317 SNPs and no plausible gene association has been made to date. We nominate rs181391313, a SNP occurring within a putative microglia-specific intronic enhancer of the Stabilin 1 (*STAB1*) gene (Figure 5a). *STAB1* is a large transmembrane receptor protein that functions in lymphocyte homing and endocytosis of ligands such as low density lipoprotein, two functions that would be consistent with a role for microglia in PD^48^. This SNP is predicted to disrupt a KLF4 binding site, consistent with the role of KLF4 in regulation of microglial gene expression^49^ (Figure 5b). Similarly, the *KCNIP3* GWAS locus resides in a 300-kb LD block harboring 94 SNPs. Our results identify two putative mediators of this phenotypic association which lead to very different functional interpretations (Figure 5c). First, rs7585473 occurs more than 250 kb upstream of the lead SNP and disrupts an oligodendrocyte-specific SOX6 motif in a peak found to interact with the Myelin and Lymphocyte (*MAL*) gene, a gene implicated in myelin biogenesis and function (Figure 5d). Alternatively, we find rs3755519 in a neuronal-specific intronic peak within the *KCNIP3* gene with clear interaction with the *KCNIP3* gene promoter. While this SNP does not show a robust machine learning prediction, nor reside within a known motif, we do identify allelic imbalance supporting its predicted functional alteration of transcription factor binding (Figure 5e). Together, these SNPs provide competing interpretations of this locus, implicating oligodendrocyte- and neuron-specific functions, and demonstrating the complexities of noncoding SNP interpretation.

**Figure 5.**
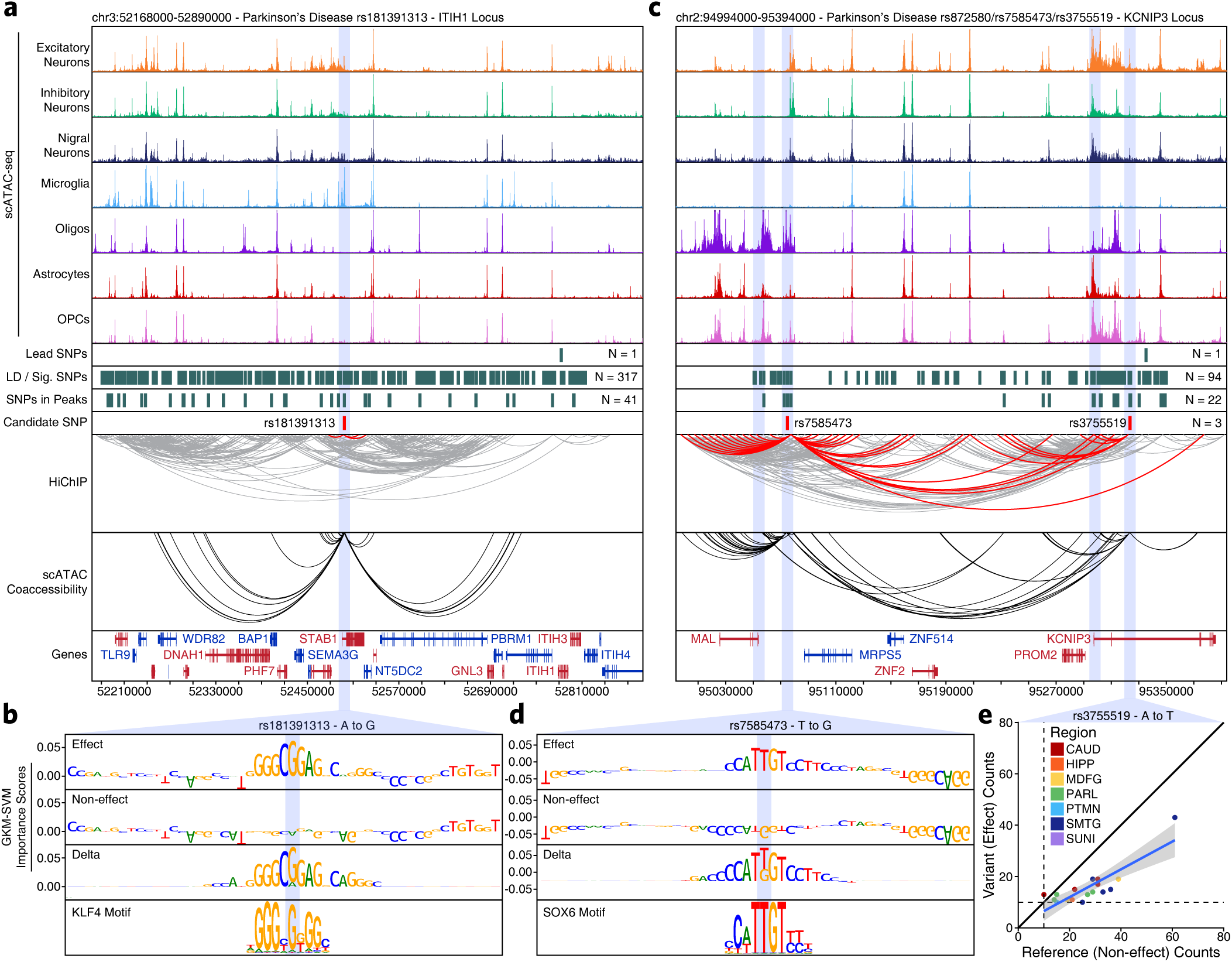
Vertical integration of multi-omic data and machine learning nominates novel gene targets in AD and PD. A. Normalized scATAC-seq-derived pseudo-bulk tracks, HiChIP loop calls, co-accessibility correlations, and machine learning predictions for LD-expanded SNPs in the *ITIH1* gene locus. For HiChIP, each line represents a loop connecting the points on each end. Red lines contain one anchor overlapping the SNP of interest while grey lines do not. B. GkmExplain importance scores for each base in the 50-bp region surrounding rs181391313 for the effect and non-effect alleles from the gkm-SVM model corresponding to microglia (Cluster 24). The predicted motif affected by the SNP is shown at the bottom and the SNP of interest is highlighted in blue. C. Sequencing tracks as shown in Figure 5a but for the *KCNIP3* locus. D. GkmExplain importance scores for each base in the 50-bp region surrounding rs7585473 for the effect and non-effect alleles from the gkm-SVM model corresponding to oligodendrocytes (Cluster 21). The predicted motif affected by the SNP is shown at the bottom and the SNP of interest is highlighted in blue. E. Dot plot showing allelic imbalance at rs3755519. The ATAC-seq counts for the reference/non-effect (A) allele and variant/effect (T) allele are shown. Each dot represents an individual bulk ATAC-seq sample colored by the brain region from which the sample was collected.

Though many such anecdotes exist (Supplementary Table 3), we also noted a pattern whereby many SNPs appear to disrupt binding sites related to the CCCTC-Binding Factor (*CTCF*) protein. For example, SNP rs6781790 disrupts a predicted CTCFL binding site within the promoter of the WD Repeat Domain 6 (*WDR6*) gene (Supplementary Fig. 9a-b). This SNP shows clear allelic imbalance across a large number of bulk ATAC-seq samples (Supplementary Fig. 9c). Similarly, SNP rs7599054 disrupts a putative CTCF binding site near the Transmembrane Protein 163 (*TMEM163*) gene (Supplementary Fig. 9d-e).

Taken together, this vertical integration of multi-omic data provides an unprecedented resolution of the landscape of inherited noncoding variation in neurodegenerative disease. Moreover, this framework and data can be applied to inform the molecular ontogeny of any brain-related GWAS polymorphism, extending the applicability of this work to all neurological disease.

### Epigenomic dissection of the *MAPT* locus explains haplotype-specific changes in local gene expression

One of the most common PD-associated risk loci is the microtubule associated protein tau (*MAPT*) gene locus. *MAPT* encodes tau proteins, a primarily neuronal set of isoforms whose pathological, hyperphosphorylated aggregates form the neurofibrillary tangles of AD^50^; however, despite the long known genetic association, it remains unclear how the *MAPT* locus may play a role in PD.

The *MAPT* locus is present within a large 1.8-Mb LD block and manifests as two distinct haplotypes, H1 and H2, which differ genetically in two primary ways: (i) more than 2000 SNPs differ across the two haplotypes, and (ii) an approximately 1-Mb inversion that includes the *MAPT* gene^51, 52^ (Figure 6a). Previous reports have nominated multiple explanations for how these alterations are associated with PD, including increased *MAPT* expression in the H1 haplotype^53, 54^ (Figure 6b), different ratios of splice isoforms^55–57^, and the use of alternative promoters^58^. We created a haplotype-specific map of chromatin accessibility and 3D chromatin interactions at the *MAPT* locus (Figure 6c). Using data from heterozygote H1/H2 individuals, we split reads into H1 and H2 haplotypes based on the presence of one of the 2366 haplotype divergent SNP (Supplementary Table 11; see methods). We tiled the region into non-overlapping 500-bp bins (to avoid biases in peak calling) and performed a Wilcoxon rank sum test to identify regions that are differentially accessible both between H1/H1 and H2/H2 homozygotes and between split reads from H1/H2 heterozygotes (Supplementary Fig. 10a-b). This identified 28 bins including an H1-specific putative enhancer 68 kb upstream of the *MAPT* promoter and the promoter of the KAT8 regulatory NSL complex subunit 1 (*KANSL1*) gene located 330 kb downstream of *MAPT* (Figure 6d (asterisks) and Supplementary Fig. 10c). Using our HiChIP data, we performed haplotype-specific virtual 4C to determine if any of these changes in chromatin accessibility were accompanied by changes in 3D chromatin interaction frequency. We identified H2-specific 3D interactions between a putative domain boundary upstream of *MAPT* (labeled “A”) and the region surrounding the *KANSL1* promoter (labeled “B”) spanning a distance of more than 600 kb inside of the inversion breakpoints (Figure 6d). Additionally, the H1-specific putative enhancer upstream of *MAPT* showed increased interaction with a second putative enhancer intronic to *MAPT* as well as with the *MAPT* promoter (Figure 6d).

**Figure 6.**
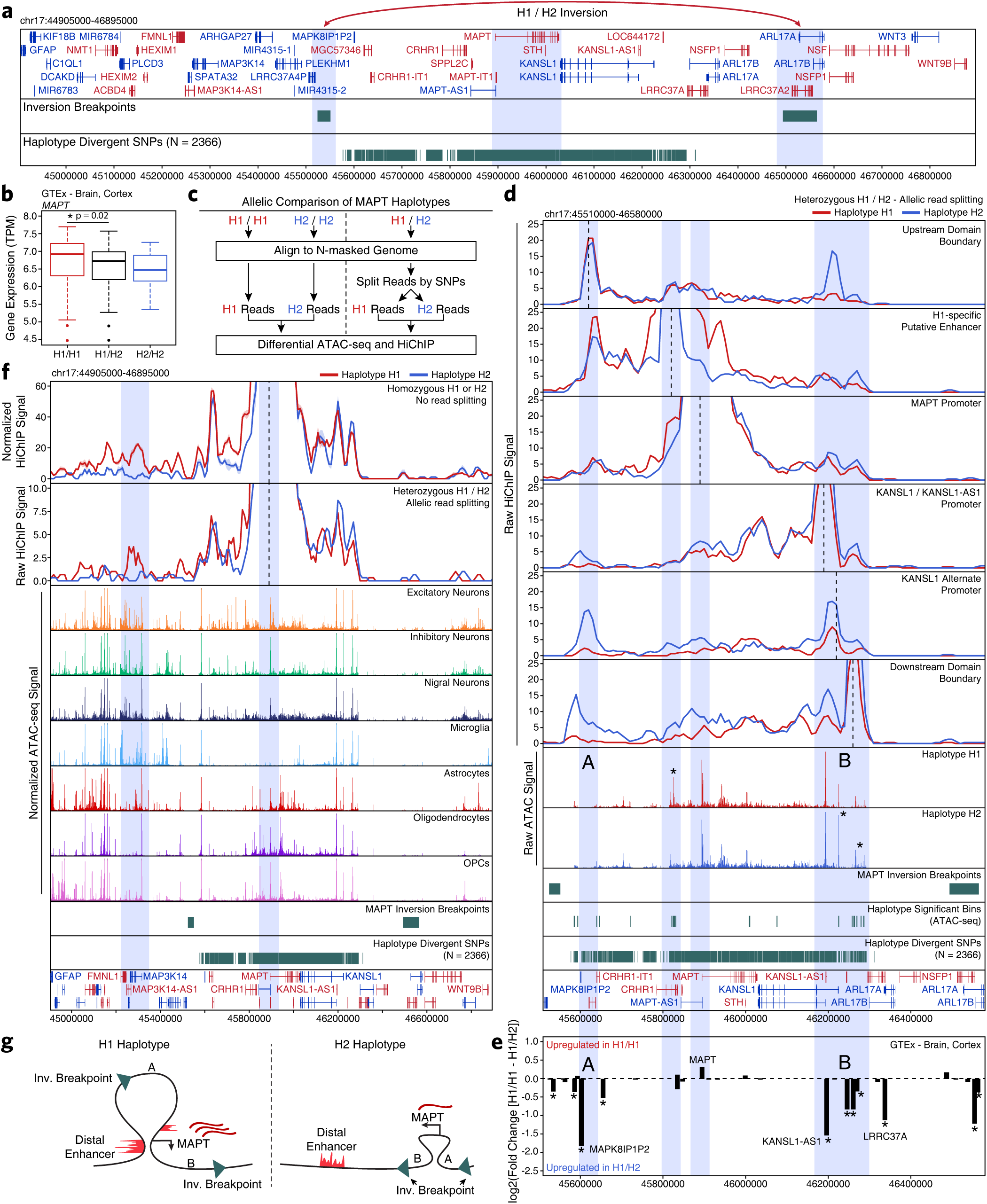
Epigenetic deconvolution of MAPT locus explains haplotype-associated transcriptional changes. A. Schematic of the *MAPT* locus (chr17:44905000-46895000) showing all genes, the predicted locations of the inversion breakpoints, and the 2366 haplotype-divergent SNPs used for haplotype-specific analyses. B. Gene expression of the *MAPT* gene shown as a box plot from GTEx cortex brain samples subdivided based on *MAPT* haplotype. The lower and upper ends of the box represent the 25th and 75th percentiles. The whiskers represent 1.5 multiplied by the inter-quartile range. C. Schematic for the allelic analysis of the *MAPT* region. Data from homozygous H1 and H2 individuals are directly compared. Data from heterozygous H1/H2 individuals are first split based off of the presence of haplotype-divergent SNPs in the reads and then compared. D. HiChIP (top) and ATAC-seq (middle) sequencing tracks of the region representing the *MAPT* locus inside of the predicted inversion breakpoints (chr17:45510000-46580000; bottom). Each track represents the merge of all available H1 or H2 reads from all heterozygotes. HiChIP and ATAC-seq tracks represent unnormalized data from heterozygotes where reads were split based on haplotype. No normalization was performed because each sample is internally controlled for allelic depth. HiChIP is shown as a virtual 4C plot where the anchor is indicated by a dotted line and the signal represents paired-end tag counts overlapping a 10-kb bin. Regions showing significant haplotype bias in ATAC-seq are marked by an asterisk. E. GTEx cortex gene expression of genes in the *MAPT* locus comparing H1 homozygotes to H1/H2. Regions A and B are shown as in Figure 6d. *p < 0.05 after multiple hypothesis correction. F. HiChIP (top) and cell type-specific scATAC-seq (middle) sequencing tracks of the region representing the *MAPT* locus outside of the predicted inversion breakpoints (bottom). HiChIP tracks for bulk homozygote H1 or H2 samples (normalized based on reads-in-loops) are shown at the top while haplotype-specific tracks from heterozygotes (unnormalized) are shown below. In each HiChIP plot, the anchor represents the *MAPT* promoter. G. Schematic illustrating the predicted haplotype-specific change in long-distance interaction between the *MAPT* promoter and the predicted distal enhancer identified in Figure 6d. Regions marked A and B represent the same regions marked in Figure 6d-e.

To better understand how these epigenetic changes impact local transcription, we used RNA-sequencing data from the Genotype-Tissue Expression (GTEx) database to identify genes that show significant haplotype-specific changes. In addition to the previously mentioned haplotype-specific differences in *MAPT* expression (Figure 6b), we also identified significant changes in the expression of genes near the largest changes in chromatin accessibility and 3D interaction (points “A” and “B”; Figure 6e). These genes include a *KANSL1* antisense transcript (*KANSL1-AS1*) and a pseudogene of the mitogen-activated protein kinase 8 interacting protein 1 (*MAPK8IP1P2*) (Supplementary Fig. 10d-e). These increases in gene expression could play a functional role in pathologic changes mediated by the different *MAPT* haplotypes or, more likely, could be a non-functional byproduct of the genomic inversion.

The above analyses help to understand how the genomic region inside of the *MAPT* inversion breakpoints differs between the H1 and H2 haplotypes; however, the inversion also changes the relative orientation of genes inside the breakpoints to enhancers and promoters outside of the breakpoints. In this way, the inversion could alter the 3D architecture of the locus and thus change which enhancers are able to communicate with the *MAPT* gene. In support of this hypothesis, we find a long-distance putative enhancer located 650 kb upstream of the *MAPT* gene that shows elevated interaction with the *MAPT* promoter specifically in the H1 haplotype (Figure 6f). We find support for this interaction both in HiChIP data from H1/H1 or H2/H2 homozygotes and from H1/H2 heterozygotes where the reads have been split based on haplotype divergent SNPs (Figure 6f). Indeed, we find multiple neuron-specific putative enhancers in this upstream region, consistent with the known neuron-specific expression of *MAPT* (Supplementary Fig. 10f), and an increase in overall 3D interaction between this upstream region and the region surrounding *MAPT* inside of the inversion breakpoints (Supplementary Fig. 10g). In total, our epigenomic dissection of the *MAPT* locus provides multiple plausible explanations for the haplotype-specific differences in *MAPT* expression and nominates multiple other genes who may exert haplotype-specific effects that are linked to differing PD phenotypes (Figure 6g).

## DISCUSSION

Here, we provide a high-resolution epigenetic characterization of the role of inherited noncoding variation in AD and PD. Our integrative multi-omic framework and machine learning classifier predicted dozens of functional SNPs, nominating gene and cellular targets for each noncoding GWAS locus. These predictions both inform well-studied disease-relevant genes, such as *BIN1* in AD, and predict novel gene-disease associations, such as *STAB1* in PD. This greatly expands our understanding of inherited variation in AD and PD and provides a roadmap for the epigenomic dissection of noncoding variation in neurodegenerative and other complex genetic diseases.

Our work initially focused on two clinically similar but pathologically distinct groups. All brain donors had been longitudinal participants in research cohorts, extensively evaluated within two years of death, and scored as high performers by neuropsychological testing (average interval between last evaluation and death was 362 days). We have shown previously that this cut off minimizes interval conversion to cognitive impairment or dementia^59^. One subset of these high performers had no or low levels of AD or PD neuropathologic change, and are labeled clinico-pathologic normal controls. Another subset of high performers showed neuropathologic changes of AD sufficient to warrant suspicion of dementia; this not common occurrence has several designations but is usually labeled resilient, meaning resilient to the clinical expression of pathologically determined AD. There is intense interest in what underlies resilience to AD because its mechanisms or adaptations may illuminate means to suppress disease expression and extend healthspan. Interestingly, our bulk ATAC-seq data showed no statistically significant differences in chromatin accessibility in any of the seven brain regions profiled for clinico-pathologic controls vs. resilience to AD. This likely indicates that the differences between these two clinical groups is minor, or potentially encoded in a rare cell type or a brain region not profiled in this work.

To inform inherited noncoding variation in neurodegenerative disease, we generated an epigenomic resource that spans the cellular and regional diversity of the adult brain. We used bulk ATAC-seq to profile seven distinct brain regions, identifying regional heterogeneity that is largely based on changes in cell type composition. To mitigate the contribution of cellular diversity to our analysis, we additionally performed scATAC-seq, profiling the chromatin accessibility of 70,631 individual cells. Cumulatively, this single-cell data identified 24 different cellular clusters which map to 7 distinct broad cell types (excitatory neurons, inhibitory neurons, nigral neurons, astrocytes, oligodendrocytes, OPCs, and microglia). Together, this resource captures the regional and cellular gene regulatory machinery that governs phenotypic expression of noncoding variation, thus allowing us to identify all polymorphisms that could putatively affect gene expression through overlap with peaks of chromatin accessibility (Tier 3). To further refine these putative functional variants, we identified the subset of polymorphisms that could be mapped to gene targets through 3D chromatin interactions or co-accessibility networks (Tier 2). Finally, we employed a machine learning approach to predict the subset of polymorphisms that would be likely to perturb transcription factor binding and validated these predictions with measurements of allelic imbalance (Tier 1). In total we implicate approximately 5 times as many genes in the phenotypic association of AD and PD and nominate functional noncoding variants for dozens of previously orphaned GWAS loci.

Through our integrative analysis, we additionally provide a comprehensive epigenetic characterization of the *MAPT* gene locus. The *MAPT* gene encodes tau isoforms, primarily neuronal microtubule binding proteins that, under pathologic conditions, can adopt an abnormal structure and extensive post translational modifications, a process called neurofibrillary degeneration, which is a hallmark of AD and other neurodegenerative diseases, but not PD^15^. Enigmatically, *MAPT* is a replicated risk locus for PD despite the absence of neurofibrillary degeneration^60, 61^. The *MAPT* locus, found on chromosome 17, represents one of the largest LD blocks in the human genome (1.8 Mb) and is present in two distinct haplotypes, H1 and H2, the latter formed by an approximately 900 kb inversion of H1 that occurred about 3 million years ago and is present mostly in Europeans^51^. Cumulatively, previous work supports *MAPT* haplotype-specific impacts on transcript amount, transcript stability, and alternative splicing in several neurodegenerative disorders^54, 56, 57^. We highlight multiple epigenetic avenues through which the *MAPT* gene is differentially regulated in the H1 and H2 haplotypes, thus explaining at least a portion of the molecular underpinnings of the observed *MAPT* GWAS association in PD.

We developed a multi-omic framework that provides a robust and comprehensive dissection of inherited variation in neurodegenerative disease. Moreover, the functional predictions made through our machine learning classifier and integrative analytical approach greatly expand our understanding of noncoding contributions to AD and PD. More broadly, this work represents a systematic approach to understand inherited variation in disease and provides an avenue towards the nomination of novel therapeutic targets that previously remained obscured by the complexity of the regulatory machinery of the noncoding genome.

## DATA AVAILABILITY

All data generated in this work is available through SRA (in progress).

## ACKNOWLEDGEMENTS

This work was supported by NIH NS062684, AG057707 (to T.M.), HG007735 (to H.Y.C.), HG009431 (to S.B.M./A.K.), and AG059918 (to M.R.C.). Sequencing data for this project was generated on an Illumina HiSeq 4000 supported in part by NIH award S10OD018220. Additional resources at the Stanford Center for Genomics and Personalized Medicine Sequencing Center were supported by NIH S10OD025212. H.Y.C. is an Investigator of the Howard Hughes Medical Institute.

## AUTHOR CONTRIBUTIONS

M.R.C., H.Y.C., and T.J.M conceived of and designed the project. M.R.C. and T.J.M. compiled the figures and wrote the manuscript with help and input from all authors. A.S. and M.R.C. performed bulk ATAC-seq data processing and analysis. M.R.C. performed all HiChIP data analysis with help from M.R.M and J.M.G. J.M.G., M.R.C., and A.S. performed all single-cell ATAC-seq data processing and analysis with supervision from W.J.G., A.K., S.B.M. and H.Y.C. M.J.G. performed GWAS locus curation, colocalization analysis, and GTEx analysis and L.F. and B.L. performed all LD score regression analysis with supervision from S.B.M. S.K. and A.S. performed all machine learning analysis with supervision from A.K. B.H.L., S.S., and M.R.C. performed all ATAC-seq, scATAC-seq, and HiChIP data generation with help from S.T.B. and M.R.M. K.S.M. curated the frozen tissue specimens used in this work.

## SUPPLEMENTARY FIGURE LEGENDS

**Supplementary Figure 1.**
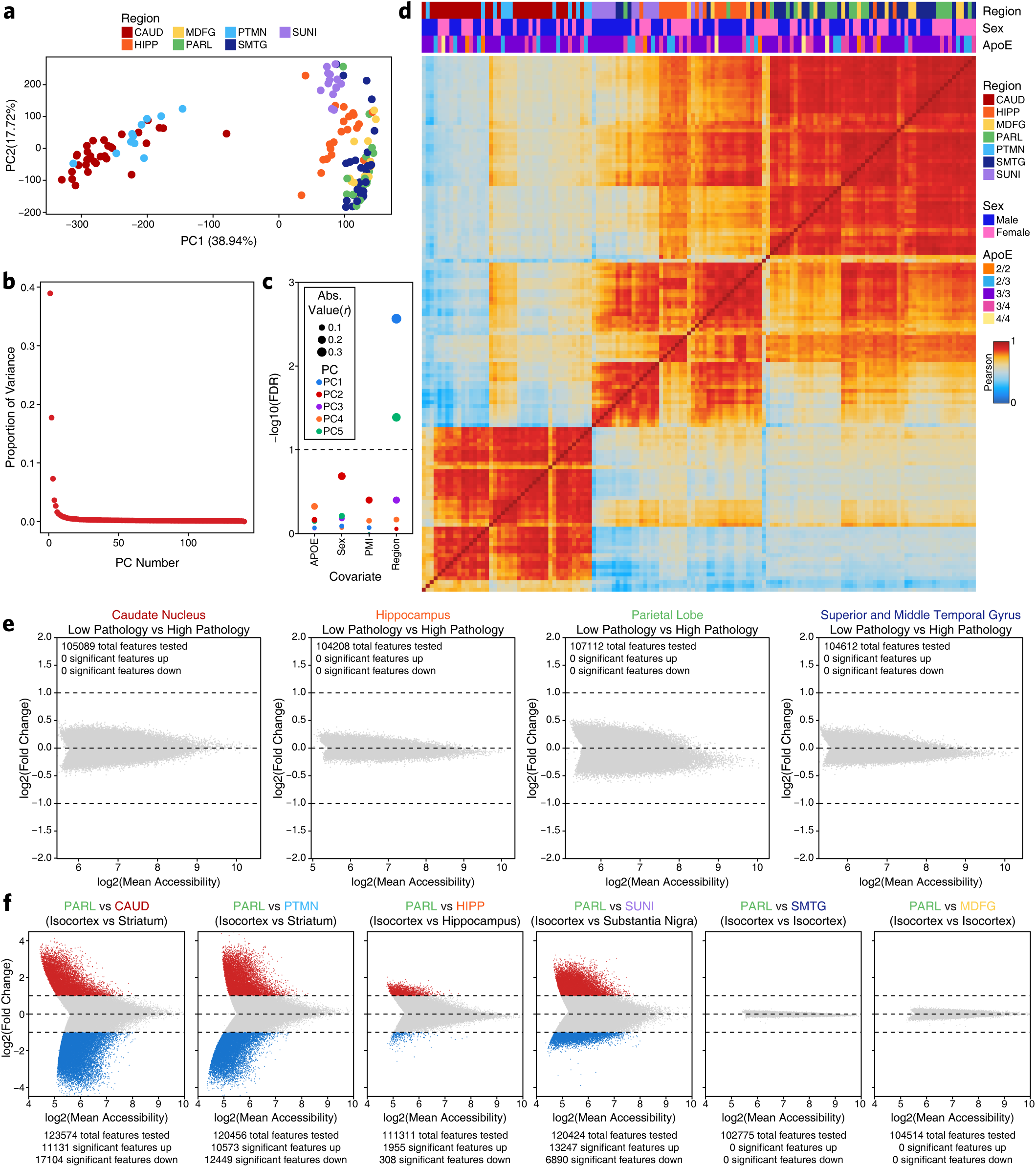
Analysis of bulk ATAC-seq data from adult brain identifies brain-regional heterogeneity. A. Principal component analysis of all samples. Each dot represents a single piece of tissue with technical replicates merged where applicable. Color represents the brain region from which the sample was isolated. B. Dot plot showing the proportion of variance explained by each principal component. C. Dot plot showing the significance of correlation between covariates and each of the top 5 principal components. Dot size represents the absolute value of the correlation while color represents the principal component number. D. Sample by sample Pearson correlation heatmap of all 140 samples profiled in this study. Brain region, donor biological sex, and *APOE* genotype are indicated colorimetrically at the top. E. MA plots showing the change in normalized bulk ATAC-seq accessibility for each peak in cognitively healthy control samples with low AD-associated pathology compared to cognitively healthy control samples with high AD-associated pathology. Each dot represents an individual peak from the merged bulk ATAC-seq peak set. Only peaks that showed non-zero accessibility in at least one sample were tested for significance. From left to right, samples from the caudate nucleus, hippocampus, parietal lobe, and superior and middle temporal gyrus are shown. F. MA plots showing the change in normalized bulk ATAC-seq accessibility comparing the parietal lobe (PARL) to all other brain regions. Each dot represents an individual peak from the merged bulk ATAC-seq peak set. Only peaks that showed non-zero accessibility in at least one sample were tested for significance.

**Supplementary Figure 2.**
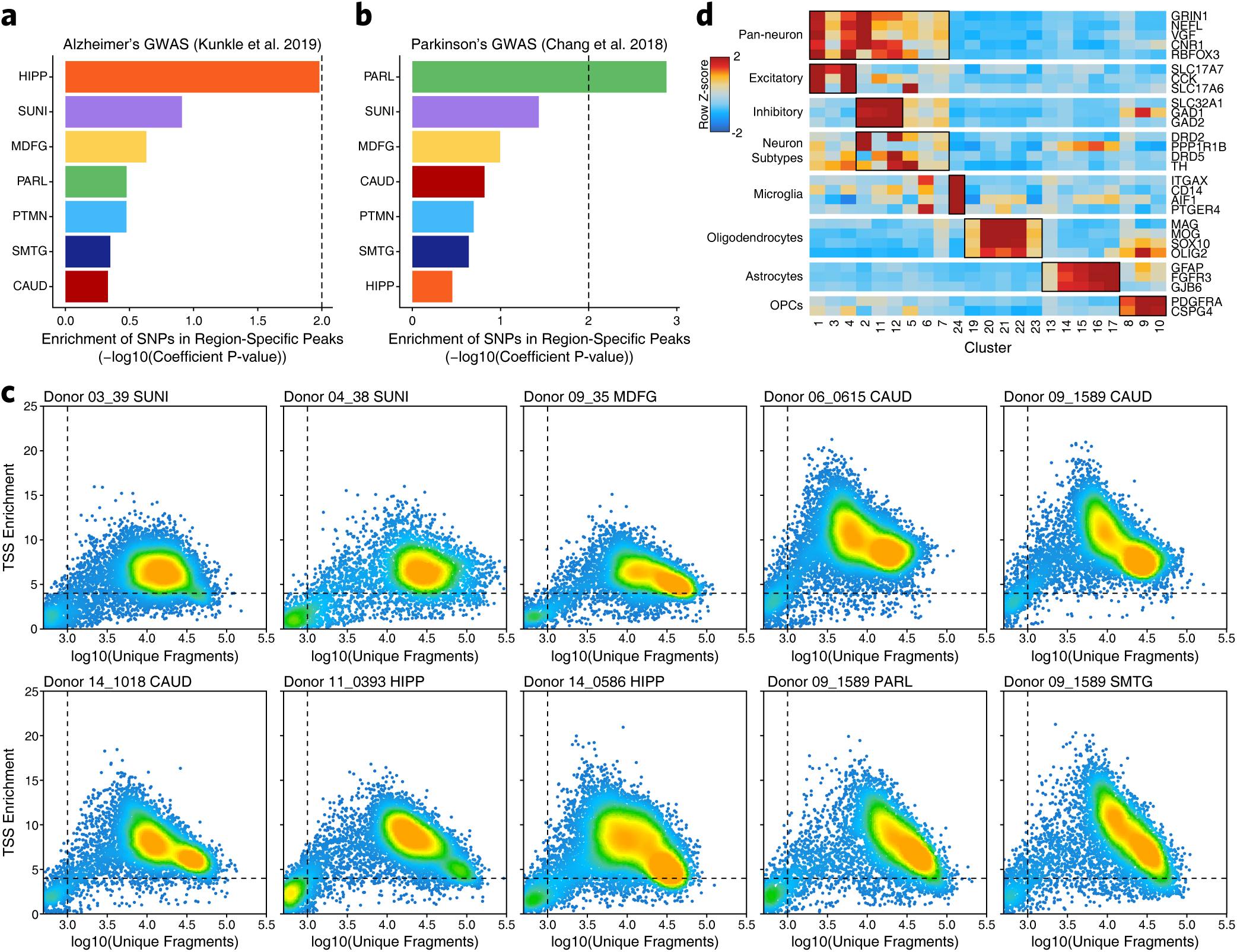
LD score regression of bulk ATAC-seq data identifies weak region-specific enrichment of AD and PD GWAS SNPs. A. Bar plot of the enrichment of AD SNPs in peaks regions of bulk ATAC-seq data from various brain regions. B. Bar plot of the enrichment of PD SNPs in peak regions of bulk ATAC-seq data from various brain regions. C. Dot plots showing the TSS enrichment score and total number of fragments for each of the 10 samples profiled by scATAC-seq. Each dot represents an individual cell. Dot color represents density on the plot. Dotted lines represent the quality control cutoffs implemented. D. Heatmap of cell type-specific markers used to identify clusters. Color represents the row-wise Z-score of chromatin accessibility in the vicinity of each gene for each cluster.

**Supplementary Figure 3.**
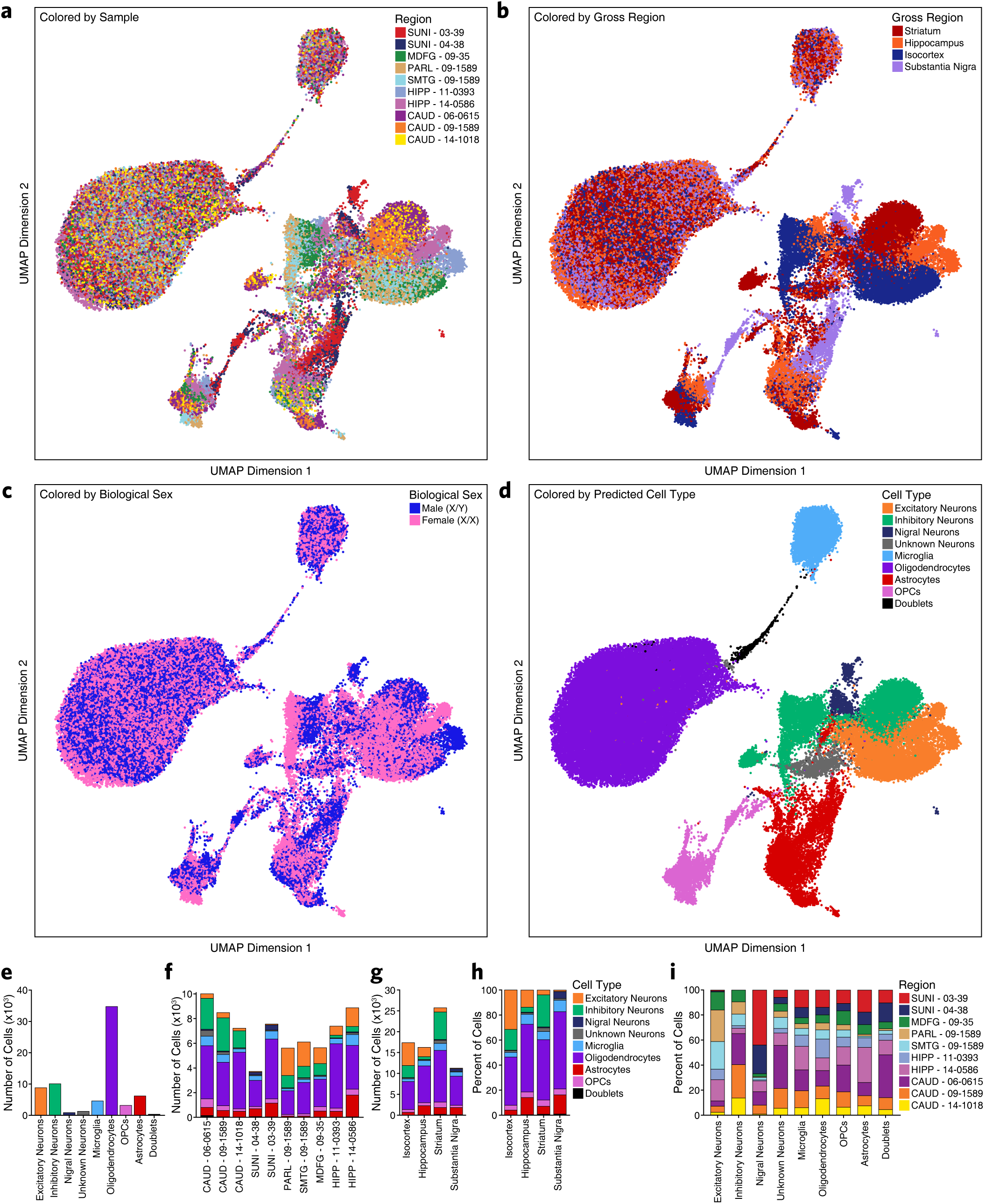
Region-centric scATAC-seq identifies cellular and regional heterogeneity in chromatin accessibility in adult brain. A. UMAP dimensionality reduction as shown in Figure 2a but colored by the sample from which each cell was generated. B. UMAP dimensionality reduction as shown in Figure 2a but colored by the brain region from which each cell was generated. C. UMAP dimensionality reduction as shown in Figure 2a but colored by the biological sex of the donor for each cell. D. UMAP dimensionality reduction as shown in Figure 2a but colored by the predicted cell type for each cell. E. Bar plot showing the number of cells identified in scATAC-seq from each of the annotated cell types. F. Bar plot showing the number of cells in scATAC-seq from each of the annotated donors/samples. Color represents the predicted cell type as shown in the legend next to Supplementary Fig. 3h. G. Bar plot showing the number of cells identified in scATAC-seq from each of the annotated cell types broken down by the brain region from which they originated. Color represents the predicted cell type as shown in the legend next to Supplementary Fig. 3h. H. Bar plot showing the percentage of each brain region composed by each cell type in scATAC-seq data. I. Bar plot showing the percentage of cells from each cell type that originated from each donor sample profiled by scATAC-seq. Color represents the biological sample from which the data was collected.

**Supplementary Figure 4.**
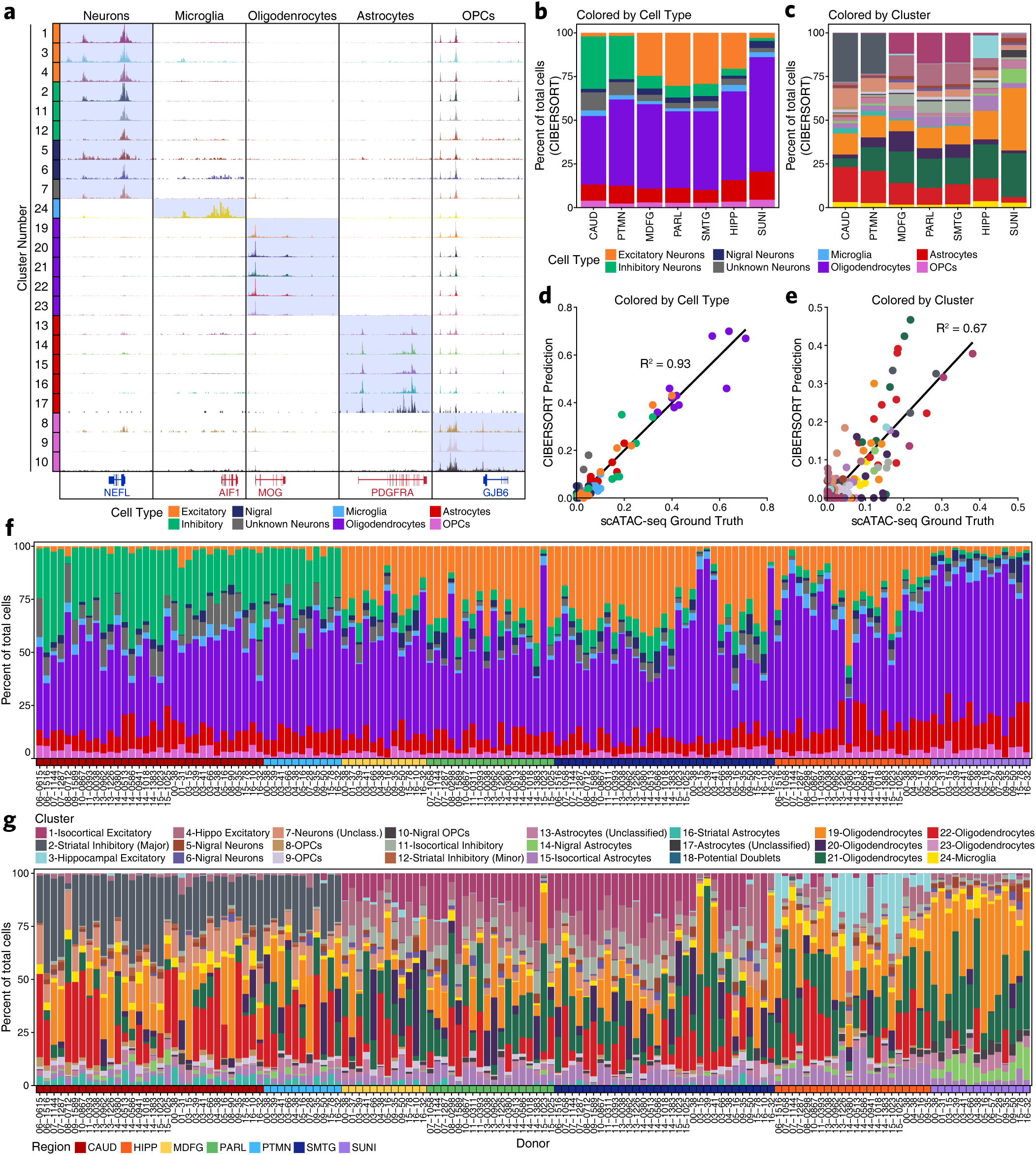
Cell type-specific scATAC-seq data enables deconvolution of chromatin accessibility data from bulk regions in the adult brain. A. Sequencing tracks of lineage-defining factors shown across all 24 scATAC-seq clusters. From left to right, *NEFL* (neurons; chr8:24933431-24966791), *AIF1* (aka *IBA1*, microglia; chr6:31607841-31617906), *MOG* (oligodendrocytes; chr6:29652183-29699713), *PDGFRA* (OPCs; chr4:54209541-54303643), and *GJB6* (astrocytes; chr13:20200243-20239571). B. Bar plot showing CIBERSORT deconvolution of bulk ATAC-seq data based on reference cell populations derived from scATAC-seq data. Clusters were subdivided into the 8 groups shown in the legend. These groups were used to preserve as much diversity as possible while merging clusters with little divergence (i.e. oligodendrocyte clusters #19-23). Bars represent the average of all bulk ATAC-seq samples profiled in the given brain regions. C. Bar plot showing CIBERSORT deconvolution of bulk ATAC-seq data based on clusters derived from scATAC-seq data. Color represents the cluster as shown in the legend of Supplementary Fig. 4g. Bars represent the average of all bulk ATAC-seq samples profiled in the given brain regions. D. Dot plot showing the performance of the CIBERSORT classifier by comparing the “ground truth” from scATAC-seq data and the CIBERSORT prediction on the bulk ATAC-seq data from the same tissue sample. Each dot represents a cell type (i.e. the merge of multiple clusters) from one of the 10 scATAC-seq samples profiled. Dots are colored by cell type according to the legend above the plot. E. Dot plot showing the performance of the CIBERSORT classifier by comparing the “ground truth” from scATAC-seq data and the CIBERSORT prediction on the bulk ATAC-seq data from the same tissue sample. Each dot represents a cluster from one of the 10 scATAC-seq samples profiled. Dots are colored by cluster according to the legend in Supplementary Fig. 4g. F. Bar plot showing CIBERSORT predictions across all bulk ATAC-seq data generated in this study. Samples are sorted and colored (bottom of plot) by the region from which they were profiled as indicated in the legend below Supplementary Fig. 4g. Bars are colored by the predicted cell type. Donor IDs are annotated below the plot. G. Bar plot showing CIBERSORT predictions across all bulk ATAC-seq data generated in this study. Samples are sorted and colored (bottom of plot) by the region from which they were profiled. Bars are colored by the predicted cluster. Donor IDs are annotated below the plot.

**Supplementary Figure 5.**
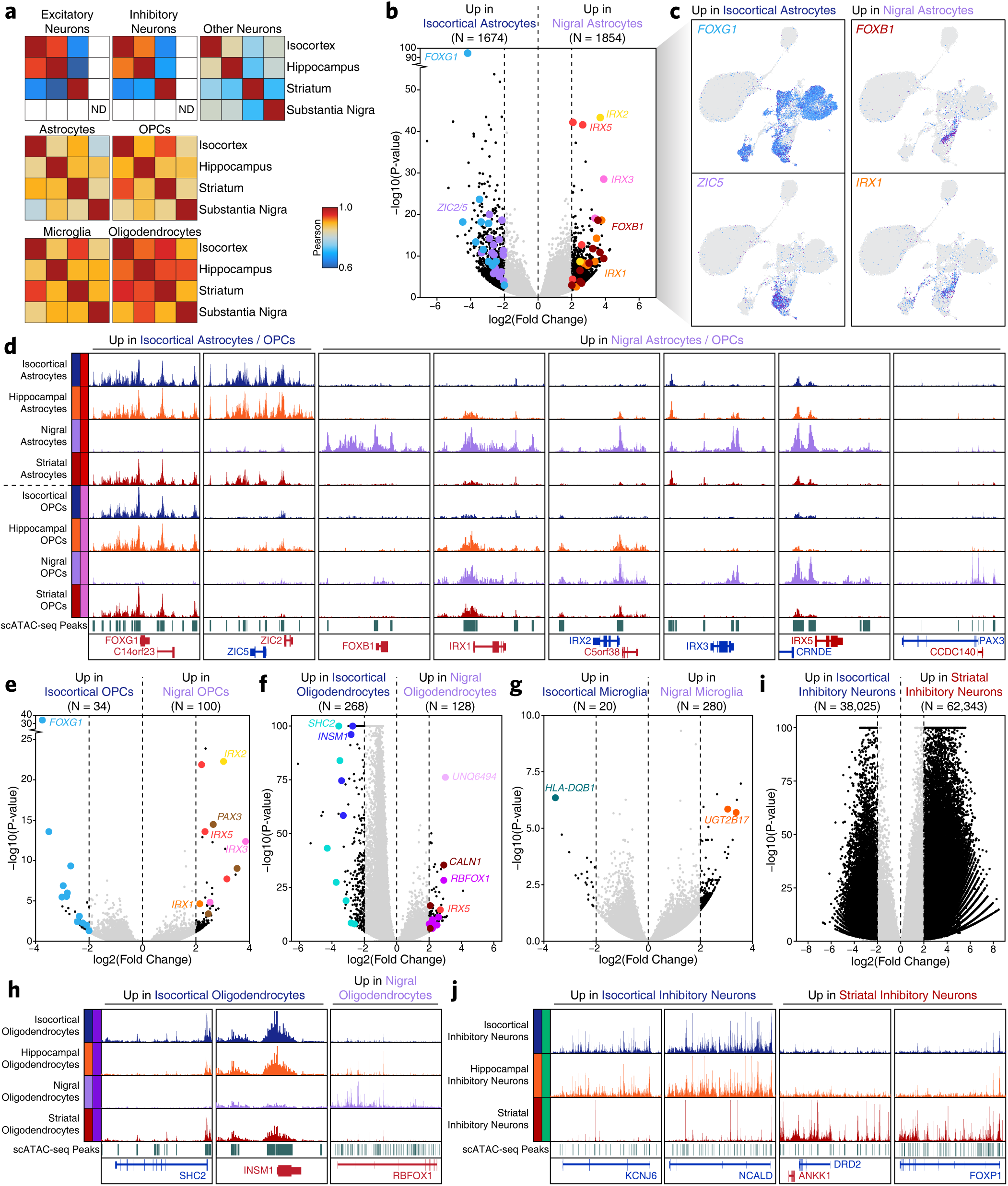
scATAC-seq reveals epigenetic encoding of region-specific cellular gene regulatory programs. A. Pearson correlation heatmaps showing the correlation of cell types across brain regions. Cell type signals were generated by making at least 2 non-overlapping pseudo-bulk replicates of at least 150 cells. Cases where insufficient cells were present to make these pseudo-bulk replicates were excluded from analysis (ND) to avoid overinterpretation. All heatmaps use the same color scale. B. Volcano plot of peaks that show differential signal between astrocytes from the substantia nigra and astrocytes from the isocortex. Peaks below a log2(fold change) threshold of 2 were not considered. Peaks near genes that are predicted to be key lineage-defining genes are accented with larger colored dots. C. UMAP dimensionality reduction plots showing gene activity scores colorimetrically for the 4 lineage-defining genes identified in Supplementary Fig. 5b (*FOXG1*, *ZIC5*, *FOXB1*, *IRX1*). D. Sequencing tracks of the multiple genomic regions showing differential chromatin accessibility between astrocytes or OPCs in the isocortex and substantia nigra. From left to right: Isocortex-specific - *FOXG1* (chr14:28750000-28787000), and *ZIC2*/*ZIC5* (chr13:99937000-99999000); Substantia Nigra-specific:-*FOXB1* (chr15:59996000-60012000), *IRX1* (chr5:3589600-3607800), *IRX2* (chr5:2737000-2760000), *IRX3* (chr16:54277000-54292000), *IRX5* (chr16:54927000-54940000), and *PAX3* (chr2:222189500-222333500). Peaks called in scATAC-seq data are shown below each plot. Sequencing tracks were derived from merging of all single cells corresponding to the annotated cell types in the specified regions. E. Volcano plot of peaks that show differential signal between OPCs from the substantia nigra and OPCs from the isocortex. Peaks below a log2(fold change) threshold of 2 were not considered. Peaks near genes that are predicted to be key lineage-defining genes are accented with larger colored dots. F. Same as Supplementary Fig. 5e but for oligodendrocytes in the substantia nigra and isocortex. G. Same as Supplementary Fig. 5e but of microglia in the substantia nigra and isocortex. H. Sequencing tracks of regions identified as differentially accessible in oligodendrocytes from the substantia nigra and isocortex. From left to right: Isocortex-specific - *SHC2* (chr19:409800-463200), and *INSM1* (chr20:20361000-20374000); Substantia nigra-specific - *RBFOX1* (chr16:5899200-7791000). Sequencing tracks were derived from merging of all single cells corresponding to the annotated cell types in the specified regions. I. Same as Supplementary Fig. 5e but for inhibitory neurons in the isocortex and striatum. J. Sequencing tracks of regions identified as differentially accessible in inhibitory neurons from the striatum and isocortex. From left to right: Isocortex-specific - *KCNJ6* (chr21:37583000-37955000), and *NCALD* (chr8:101673000-102141000); Striatum-specific - *DRD2* (chr11:113369000-113602000), and *FOXP1* (chr3:70922000-71622000).Sequencing tracks were derived from merging of all single cells corresponding to the annotated cell types in the specified regions.

**Supplementary Figure 6.**
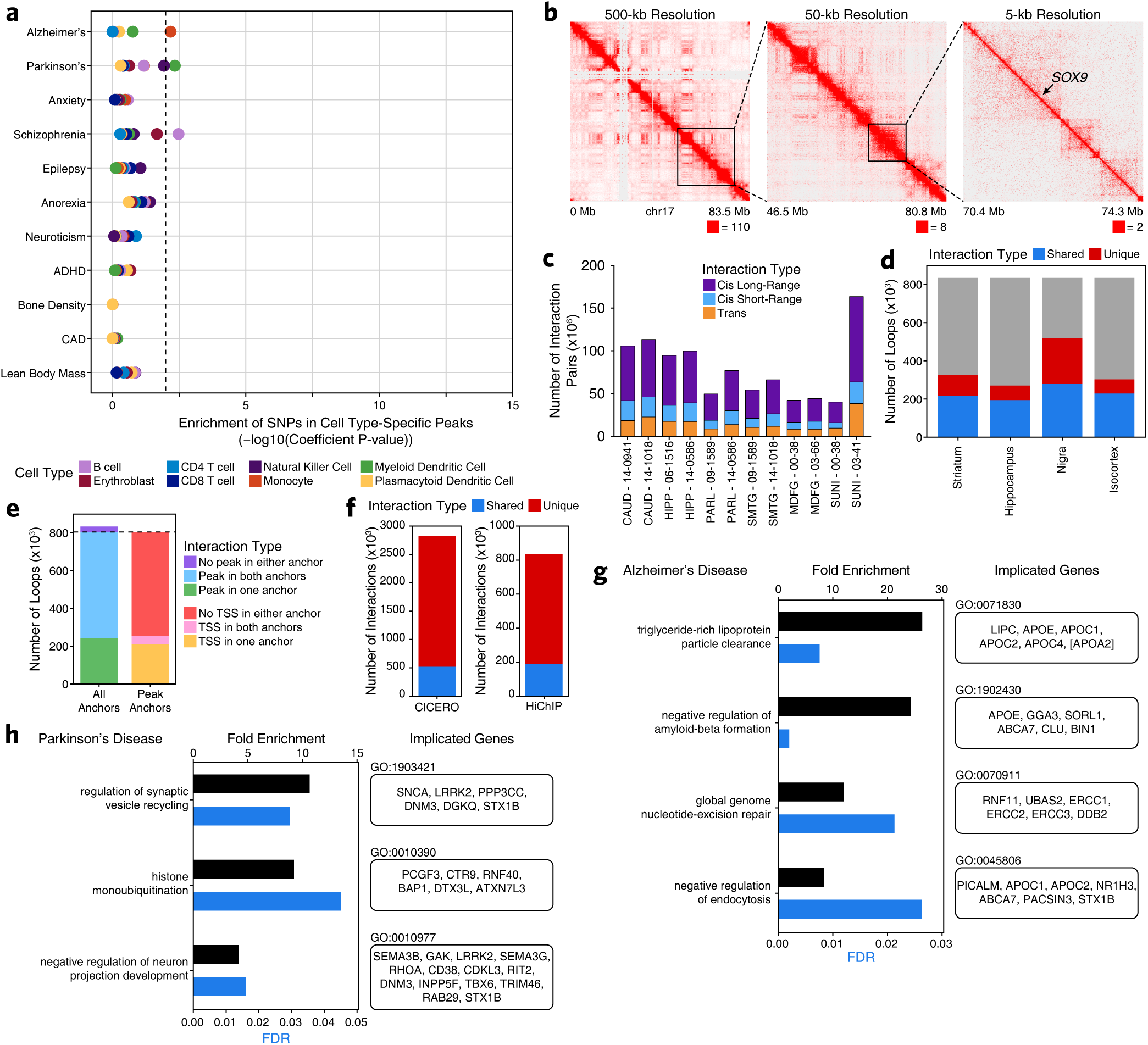
HiChIP implicates disease-relevant genes in AD and PD through linkage of noncoding GWAS SNPs to target genes. A. LD score regression identifying the enrichment of GWAS SNPs from various brain- and non-brain-related conditions in the peak regions of bulk ATAC-seq data from various hematopoietic cell types as indicated by color. B. Heatmap representation of HiChIP interaction signal at 100-kb, 25-kb, and 5-kb resolution at the SOX9 locus. C. Bar plots showing the number of valid interaction pairs identified in HiChIP data from all samples profiled in this study. Color represents the type of interaction identified. D. Bar plot showing the overlap of FitHiChIP loop calls from the 4 gross brain regions profiled. Color indicates whether the loop was identified in a single region (unique) or more than one region (shared). E. Bar plot showing the classification of FitHiChIP loop calls based on whether the loop call contained an ATAC-seq peak (bulk or single-cell) or TSS in one, both, or no anchor. F. Bar plots showing the number of Cicero-predicted co-accessibility-based peak links that are observed in HiChIP (left) or the number of HiChIP-based FitHiChIP loop calls that are predicted as peak links by Cicero. G. GO-term enrichments of genes linked to AD GWAS SNPs. H. GO-term enrichments of genes linked to PD GWAS SNPs.

**Supplementary Figure 7.**
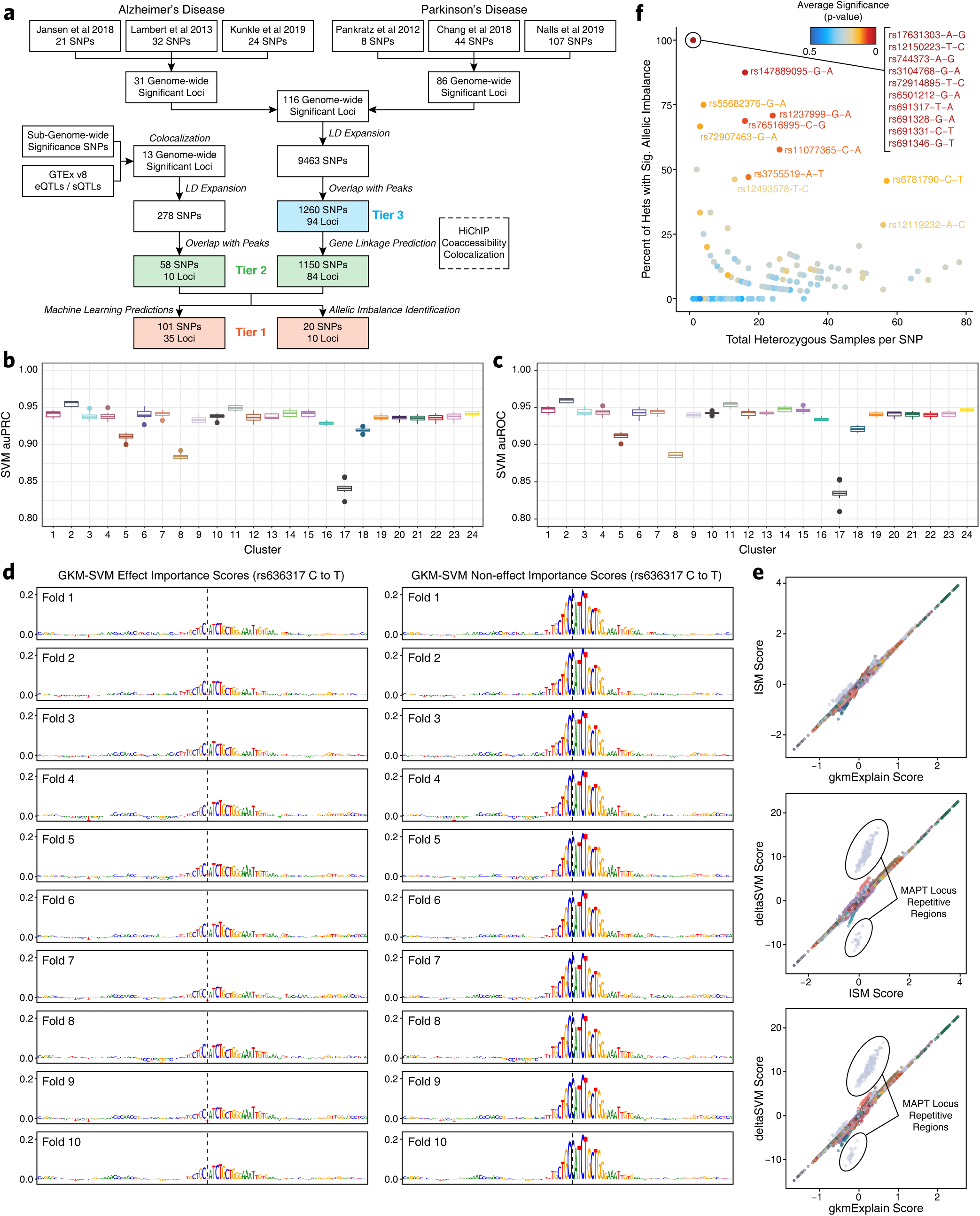
Machine learning and allelic imbalance predict functional noncoding SNPs in AD and PD. A. Flow chart of the analytical framework used to prioritize noncoding SNPs and predict functionality. The highest confidence SNPs (Tier 1) are supported by either machine learning predictions, allelic imbalance, or both. Moderate confidence SNPs (Tier 2) are supported by the presence of the SNP within a peak and a HiChIP loop or co-accessibility peak link that connects the SNP to a gene. Lower confidence SNPs (Tier 3) are only supported by the presence of the SNP in a peak. B. Box plot showing the area under the precision-recall curve for the gkm-SVM machine learning classifier. Performance for each cluster is shown with dots representing outliers. The lower and upper ends of the box represent the 25th and 75th percentiles. The whiskers represent 1.5 multiplied by the inter-quartile range. C. Box plot showing the area under the receiver-operating characteristics curve for the gkm-SVM machine learning classifier. Performance for each cluster is shown with dots representing outliers. The lower and upper ends of the box represent the 25th and 75th percentiles. The whiskers represent 1.5 multiplied by the inter-quartile range. D. GkmExplain importance scores shown across all 10 folds for each base across a 100-bp window surrounding rs636317 for the effect (left) and noneffect (right) bases. E. Dot plots showing comparison of the GkmExplain score, ISM score, and deltaSVM score. Each dot represents an individual SNP test in a given fold. Dot color represents the GWAS locus number. The only off-diagonal dots (circled) correspond to repetitive regions within the *MAPT* locus where the deltaSVM score appears to be particularly sensitive. F. Dot plot showing allelic imbalance across all bulk ATAC-seq data used in this study. ATAC-seq data was used to genotype individuals to identify heterozygotes. Allelic imbalance was defined as ratio of wildtype to variant reads that passes the binomial test with a p-value less than 0.05. Color indicates the average significance of the binomial test across all heterozygotes.

**Supplementary Figure 8.**
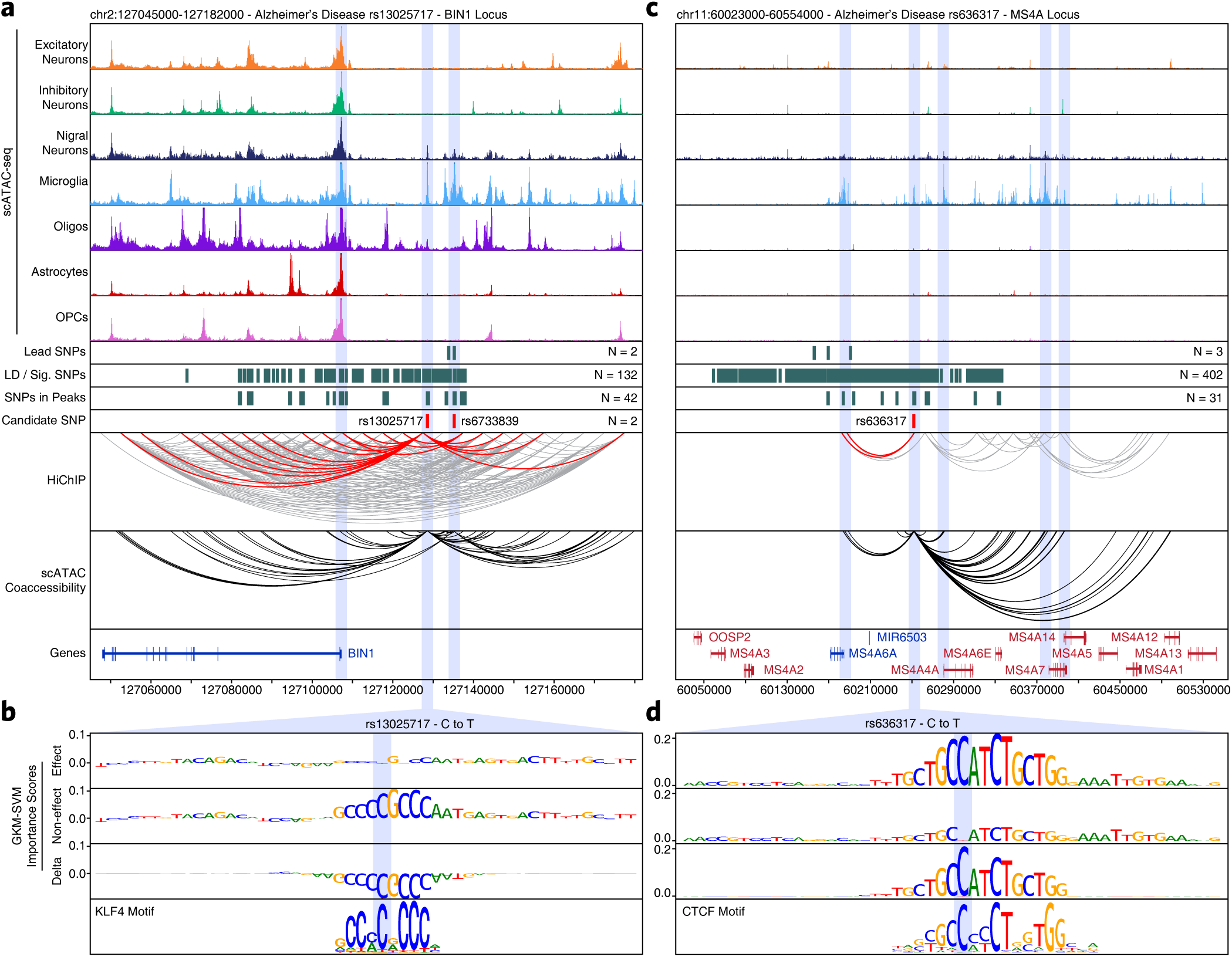
Multi-omic characterization of well-studied AD-related GWAS loci pinpoints putative functional noncoding SNPs. A. Normalized scATAC-seq-derived pseudo-bulk tracks, HiChIP loop calls, co-accessibility correlations, and machine learning predictions for LD-expanded SNPs in the *BIN1* locus. For HiChIP, each line represents a loop connecting the points on each end. Red lines contain one anchor overlapping the SNP of interest while grey lines do not. B. GkmExplain importance scores for each base in the 50-bp region surrounding rs13025717 for the effect and non-effect alleles from the gkm-SVM model for microglia (Cluster 24). The predicted motif affected by the SNP is shown at the bottom and the SNP of interest is highlighted in blue. C. Sequencing tracks as shown in Supplementary Fig. 8a but for the *MS4A* gene locus. D. GkmExplain importance scores for each base in the 50-bp region surrounding rs636317 for the effect and non-effect alleles from the gkm-SVM model for microglia (Cluster 24). The predicted motif affected by the SNP is shown at the bottom and the SNP of interest is highlighted in blue.

**Supplementary Figure 9.**
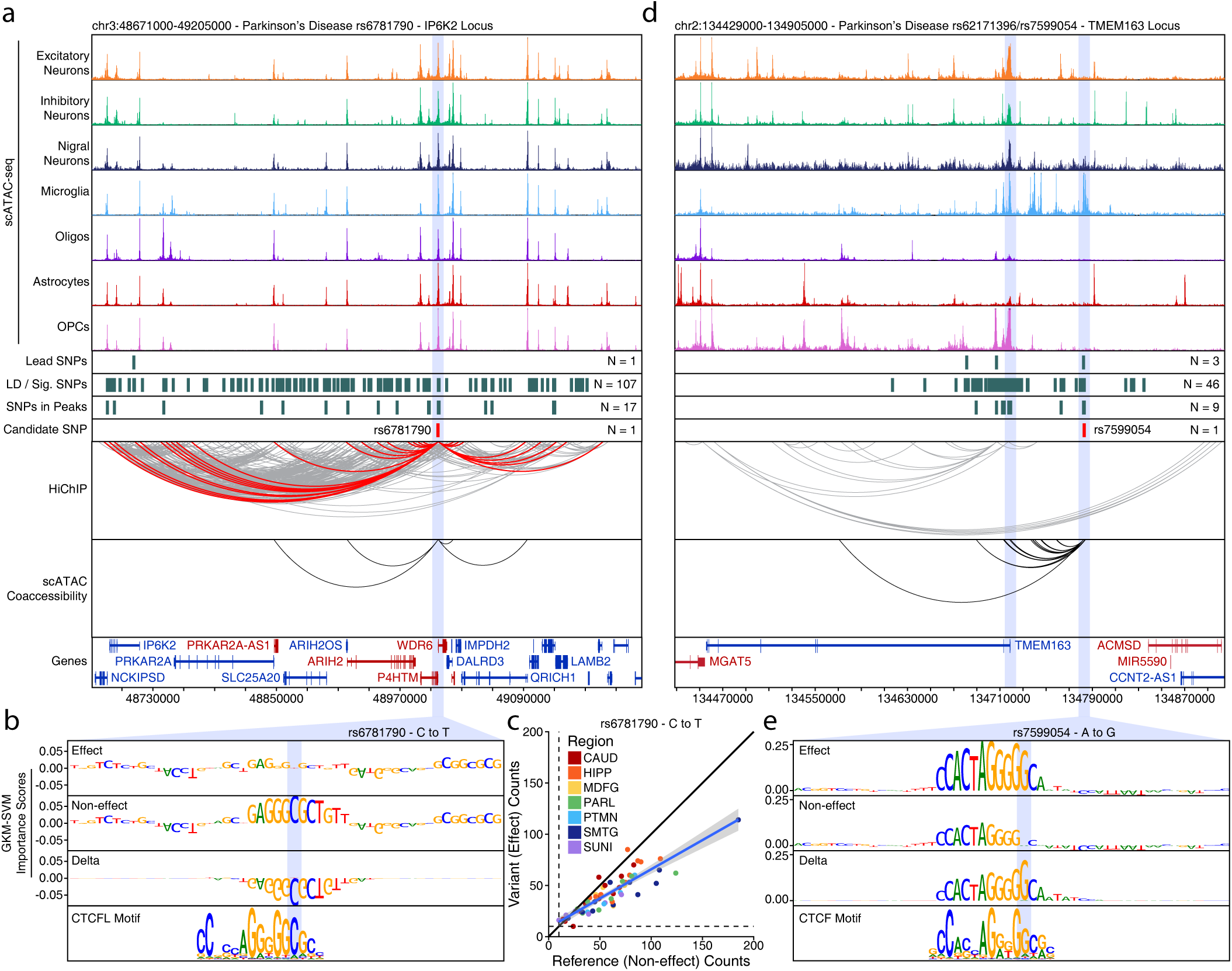
Multi-omic characterization of noncoding SNPs identifies novel genes implicated in PD. A. Normalized scATAC-seq-derived pseudo-bulk tracks, HiChIP loop calls, co-accessibility correlations, and machine learning predictions for LD-expanded SNPs in the *IP6K2* locus. For HiChIP, each line represents a loop connecting the points on each end. Red lines contain one anchor overlapping the SNP of interest while grey lines do not. B. GkmExplain importance scores for each base in the 50-bp region surrounding rs6781790 for the effect and non-effect alleles from the gkm-SVM model for astrocytes (Cluster 15). The predicted motif affected by the SNP is shown at the bottom and the SNP of interest is highlighted in blue. C. Dot plot showing allelic imbalance at rs6781790. The ATAC-seq counts for the reference/non-effect (C) allele and variant/effect (T) allele are plotted. Each dot represents an individual bulk ATAC-seq sample colored by the brain region from which the sample was collected. D. Sequencing tracks as shown in Supplementary Fig. 9a but for the *TMEM163* locus. E. GkmExplain importance scores for each base in the 50-bp region surrounding rs7599054 for the effect and non-effect alleles from the gkm-SVM model for microglia (Cluster 24). The predicted motif affected by the SNP is shown at the bottom and the SNP of interest is highlighted in blue.

**Supplementary Figure 10.**
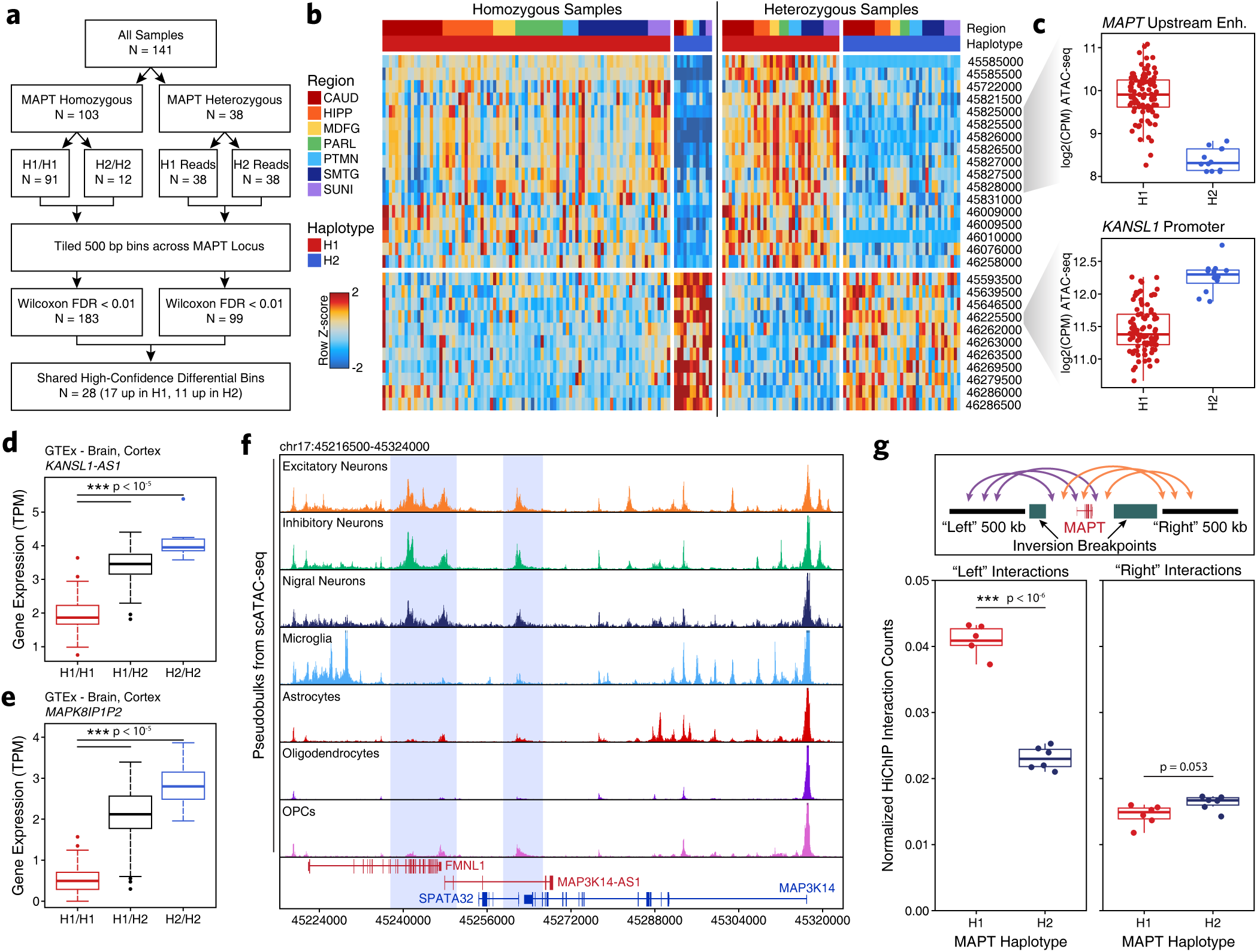
Epigenomic dissection of the MAPT locus. A. Flowchart illustrating the analytical scheme used to identify bins with significant allelic imbalance across the H1 and H2 *MAPT* haplotypes. B. Heatmaps showing chromatin accessibility in 500-bp bins identified as having significantly different accessibility across *MAPT* haplotypes. Regions are shown for homozygous samples without allelic read splitting (left) and for heterozygous samples after allelic read splitting (right). Bin start coordinates are shown to the right. C. Box and whiskers plots for multiple regions which show differential chromatin accessibility across the H1 and H2 *MAPT* haplotypes. Each dot represents a single homozygous H1 or homozygous H2 sample. Heterozygotes are not shown. The lower and upper ends of the box represent the 25th and 75th percentiles. The whiskers represent 1.5 multiplied by the inter-quartile range. D. Gene expression of the *KANSL1-AS1* gene shown as a box plot from GTEx cortex brain samples subdivided based on *MAPT* haplotype. The lower and upper ends of the box represent the 25th and 75th percentiles. The whiskers represent 1.5 multiplied by the inter-quartile range. ***p < 10^-5^. E. Gene expression of the *MAPK8IP1P2* gene shown as a box plot from GTEx cortex brain samples subdivided based on *MAPT* haplotype. The lower and upper ends of the box represent the 25th and 75th percentiles. The whiskers represent 1.5 multiplied by the inter-quartile range. ***p < 10^-5^. F. Sequencing tracks from pseudo-bulk data derived from predicted cell types in scATAC-seq data. This region represents a zoomed in view of the predicted distal enhancer region (chr17:45216500-45324000) that interacts with the *MAPT* promoter in the H1 haplotype. Putative neuron-specific enhancers are highlighted in blue. G. Box plots showing differential HiChIP interaction signal occurring between regions within the *MAPT* inversion and regions outside the inversion (“left” or “right”). The schematic at the top explains the analysis performed. The box plots show normalized HiChIP interaction counts for the H1 and H2 haplotypes for upstream/“left” interactions and downstream/“right” interactions.

## SUPPLEMENTARY TABLES

**Supplementary Table 1** – Donor information and sequencing statistics for all samples profiled by bulk ATAC-seq, scATAC, and HiChIP.

**Supplementary Table 2** – Final merged peak set derived from all bulk ATAC-seq data.

**Supplementary Table 3** – All LD-expanded GWAS SNPs from AD and PD and their relevant metadata and characterizations.

**Supplementary Table 4** – Quality control information for all individual cells profiled by scATAC-seq and the cluster residence information for all clusters and samples.

**Supplementary Table 5** – Final merged peak set derived from all scATAC-seq data.

**Supplementary Table 6** – Results of feature binarization from scATAC-seq data showing cell type-specific peaks.

**Supplementary Table 7** – CIBERSORT signature matrices for the cell group-specific and cluster-specific classifiers.

**Supplementary Table 8** – Results of differential accessibility comparisons between the substantia nigra and isocortex for astrocytes, OPCs, oligodendrocytes, and microglia.

**Supplementary Table 9** – Results of all LD score regression analyses across all conditions and cell types.

**Supplementary Table 10** – All FitHiChIP loop calls overlapping a SNP on at least one anchor.

**Supplementary Table 11** – All SNPs that are divergent between the H1 and H2 haplotypes in the *MAPT* locus.

## METHODS

### Code Availability

All custom code used in this work is available in the following GitHub repository: https://github.com/kundajelab/alzheimers_parkinsons.

### Publicly Available Data Used In This Work

All QTL analysis was performed using GTEx v8. Additionally, we downloaded full-genome summary statistics of GWAS associations for three Alzheimer’s cohorts^1–3^ and three Parkinson’s cohorts^6, 7, 62^; however, it should be noted that these cohorts are not all mutually exclusive.

### Genome Annotations

All data is aligned and annotated to the hg38 reference genome.

### Sequencing

Bulk ATAC-seq, and HiChIP were sequenced using an Illumina HiSeq 4000 with paired-end 75-bp reads. Single-cell ATAC-seq was sequenced using an Illumina NovaSeq 6000 with an S4 flow cell with paired-end 99 bp reads.

### Sample acquisition and patient consent

Primary brain samples were acquired post-mortem with IRB-approved informed consent. Human donor sample sizes were chosen to provide sufficient confidence to validate methodological conclusions. Human brain samples were collected with an average post-mortem interval of 3.9 hours (range 2.0 – 6.9 hours). Macrodissected brain regions were flash frozen in liquid nitrogen. Some samples were embedded in Optimal Cutting Temperature (OCT) compound. All samples were stored at −80°C until use.

### Isolation of nuclei from frozen tissue chunks

Nuclei were isolated from frozen tissue as described previously^63, 64^. This protocol is now available on protocols.io (dx.doi.org/10.17504/protocols.io.6t8herw). After isolation, nuclei were cryopreserved in BAM Banker (Wako Chemicals) and stored at −80°C for use in other assays such as scATAC-seq and HiChIP.

### Statistics

All statistical tests performed are included in the figure legends or methods where relevant.

### ATAC-seq Data Processing

The ENCODE DCC ATAC-seq pipeline (doi:10.5281/zenodo.211733) (V1.1.7) was used to process bulk ATAC-seq samples, starting from fastq files. The pipeline was executed with IDR enabled and the IDR threshold set to 0.05. The GRCh38 reference genome assembly was used, keeping only the primary chromosomes chr1 - chr22, chrX, chrY, chrM. The pipeline was executed with ATAQC enabled, using GENCODE version 29 TSS annotations. Biological replicates were analyzed individually, with the two technical replicates for each bio-rep provided as inputs to the “atac.bams” argument of the pipeline. Other arguments to the pipeline were kept at their defaults.

### ATAC-seq Peak Calling

Pipeline peak calls underwent several levels of filtering to identify credible peak sets. The IDR optimal peak set from the DCC pipeline for each biological replicate was determined. It was observed that although the IDR peaks for individual biological replicates were corrected for multiple testing, the high number of biological samples in the dataset served as another source of multiple testing error. To address this source of error, tagAlign files for all biological replicates for a given brain region/ condition were concatenated. The DCC pipeline (v1.1.7) was subsequently executed on the merged tagAlign files as single-replicate inputs. The pipeline generated pseudo-replicates from the input tagAlign files for each brain region/condition. Optimal IDR peaks were called from the pseudo-replicates. This set of IDR peaks was filtered to keep peaks supported by 30 percent or more of IDR peaks from the pipeline runs on individual biological replicates.

Sample-by-peak count matrices were then generated from the resulting set of filtered peaks. Filtered peaks from the pooled tagAlign files were concatenated and truncated to within 200 base pairs of the summit (100 base pair flank kept upstream and downstream of the peak summit). These 200 bp regions were merged with the bedtools^65^ merge command to avoid merging peaks with low levels of overlap. The bedtools coverage-counts was used to compute the number of tagAlign reads that overlapped each peak region in the pseudo-replicates in the merged tagAlign dataset. This analysis yielded a total of n=186,559 peaks combined across the brain regions.

### Motif enrichment

Motif enrichment was performed using the hypergeometric test as described previously^64, 66^.

### Feature Binarization

Identification of “unique” peaks from ATAC-seq data was performed as described previously^12, 64^.

### Sequencing Tracks

Sequencing tracks were created using the WashU Epigenome Browser. All sequencing tracks of a given locus have the same y-axis. All tracks show data that has been normalized by “reads-in-peaks” (for ATAC-seq) or “reads-in-loops” for HiChIP to account for differences in signal-to-background ratios across multiple samples, unless otherwise stated. For all sequencing tracks, genes that are on the plus strand (i.e. 5’ to 3’ in the left to right direction) are shown in red and genes that are on the minus strand (i.e. 5’ to 3’ in the right to left direction) are shown in blue to enable identification of the TSS.

### LD score regression

We apply stratified LD score regression, a method for partitioning heritability from GWAS summary statistics, to sets of tissue or cell type specific ATAC-seq peaks to identify disease-relevant tissues and cell types across for Alzheimer’s and Parkinson’s diseases along with other brain-related GWAS traits. We used both bulk ATAC-seq and single cell ATAC-seq data. For bulk ATAC-seq we kept only peaks replicating in at least 30% of samples for each tissue part. ATAC-seq peaks were converted from hg38 to hg19 for analysis with GWAS data. We followed the LD score regression tutorial (https://github.com/bulik/ldsc/wiki) as used previously^67^ for bulk data and as recently developed for single-cell specific analysis^68^. We used brain related GWAS summary statistics such as Alzheimer’s^1^, Parkinson’s^6^, Schizophrenia^69^, Anorexia Nervosa^70^, Attention Deficit Hyperactivity Disorder (ADHD)^71^, Anxiety^72^, Neuroticism^73^ and Epilepsy^74^. To serve as controls, we also used summary statistics for GWAS of traits not obviously linked to brain tissues such as Lean Body Mass^75^, Bone Mineral Density^76^ and Coronary Artery Disease^77^. In particular, we looked at the regression coefficient p-value, indicative of the contribution of this annotation to trait heritability, conditional on the other annotations.

### Allelic imbalance from ATAC-seq data

Samples were first re-aligned to an N-masked version of the hg38 genome where all relevant SNP positions were changed to “N” to prevent mapping bias. Allelic depth at each desired position was obtained using samtools mpileup (v1.5) followed by varscan mpileup2snp (v2.4.3). Allele counts for the reference and variant alleles were extracted and compared using the binomial test to identify significant allelic imbalance.

### SNP selection for colocalization testing

A single test for colocalization of GWAS and eQTL association signals involves a locus, a GWAS, an eQTL tissue, and a gene expressed in that tissue. For each GWAS, we selected the set of all loci for which the lead GWAS variant had p-value < 1e-5. Using eQTLs from GTEx brain tissues in the GTEx v8 dataset, we then found all tissue-gene combinations for which the lead SNP at one of the GWAS loci had an eQTL SNP (association p-value < 1e-5) for that gene in that GTEx tissue. This resulted in a list of unique combinations of GWAS trait / genomic locus / eQTL tissue / eQTL gene, each to be tested individually for colocalization of GWAS and eQTL signals. The GWAS threshold of 1e-5 is less stringent than the threshold for genome-wide significance, but we favored sensitivity over specificity when selecting which SNPs to test, since colocalization with a strong eQTL signal may still suggest that a sub-threshold GWAS locus has an expression-mediated effect on disease.

### Colocalization analysis

For each colocalization test combination as defined above, we selected all 1000 Genomes Phase 3 variants within a window of 500kb around the lead GWAS variant. We narrowed this list down to SNPs measured not only in the 1000 Genomes VCF, but also in the GWAS and eQTL summary statistics for the selected trait, tissue, and gene. We used a streamlined version of the FINEMAP tool^78^ to compute posterior causal probabilities for each SNP at the locus in both the GWAS and eQTL studies, and then combined these probabilities as described in eCAVIAR^79^ to compute a colocalization posterior probability (CLPP) score for this test locus. We considered a SNP weakly colocalized if its CLPP score exceeded 0.01 and strongly colocalized if its CLPP score exceeded 0.05; although these seem like quite low probabilities, we have seen previously that loci exceeding this latter cutoff show strong likelihood of sharing causal variants^80^.

### Selection of candidate SNPs for ATAC-seq overlap analysis, HiChIP interaction tests, and gkm-SVM model-based allelic effect scores

Our goal was to identify SNPs with a causal effect on any of the selected GWAS traits. To minimize the chances of excluding causal GWAS SNPs, we selected the set of all variants achieving a genome-wide significant p-value < 5e-8 for any GWAS trait. We then added in any lead SNPs from the colocalization analysis that achieved CLPP score of > 0.01, even those that did not pass the genome-wide significance value of p < 5e-8. We also included all trait-associated SNPs curated from two other Parkinson’s studies^6, 7^. In these studies, full summary statistics were not publicly available for the entire genome because meta-analysis was applied only to the subset of SNPs reaching genome-wide significance in a previous Parkinson’s GWAS. We then computed the full set of SNPs that had LD R^2 ≥ 0.8 with at least one of the SNPs in the set selected above. Together, these LD buddies plus the original set of trait-relevant SNPs comprised the set of SNPs tested in our subsequent functional analyses.

### Testing GWAS loci for overlap with ATAC-seq peaks

We tested all SNPs in the above set for overlap with ATAC-seq peaks from two different annotation formats. The first annotation consisted of bulk ATAC-seq peaks identified in one of 7 brain regions. The second annotation consisted of cluster-specific peaks from single-cell ATAC-seq data. For each variant selected for functional analysis, we determined all cellular contexts in which an ATAC-seq peak contained this variant, as well as the nearest peak if no peak contained the variant.

### Single-cell ATAC-seq library generation

Cryopreserved nuclei were thawed on ice and 65,000 nuclei were transferred to a tube containing 1 ml of RSB-T [10 mM Tris-HCl pH 7.5, 10 mM NaCl, 3 mM MgCl2, 0.1% Tween]. Nuclei were pelleted at 500 RCF for 5 minutes at 4°C in a fixed angle rotor. The supernatant was fully removed using two pipetting steps (p1000 to remove down to the last 100 ul, then p200 to remove all remaining supernatant). This pellet was then gently resuspended in 12 ul of 1x Nuclei Buffer (10x Genomics). To transpose, 5 ul of this nuclei suspension (containing 27,000 nuclei) was transferred to a tube containing 10 ul of transposition mix (10x Genomics). This reaction mixture was incubated at 37°C for 1 hour to transpose. The remainder of library generation was completed as described in the 10x Genomics Single Cell ATAC Regent Kits User Guide (v1 Chemistry).

### Single-cell ATAC-seq LSI clustering and visualization

To cluster our scATAC-seq data, we first identified a robust set of peak regions followed by iterative LSI clustering^12, 18^. Briefly, we created 1-kb windows tiled across the genome and determined whether each cell was accessible within each window (binary). Next, we identified the top 50,000 accessible windows across all samples (accounting for GC bias) and performed an LSI dimensionality reduction (TF-IDF transformation followed by Singular Value Decomposition SVD) on these windows followed by Harmony batch correction^81^. We then performed Seurat^82^ clustering (FindClusters v2.3) on the harmonized LSI dimensions at a resolution of 0.8, 0.4 and 0.2, keeping the clustering for which the minimum cluster size was greater than 100 cells (0.2 if this condition is not met). For each cluster, we called peaks on the Tn5-corrected insertions (each end of the Tn5-corrected fragments) using the MACS2 callpeak command with parameters ‘--shift −75 --extsize 150 --nomodel --call-summits --nolambda --keep-dup all -q 0.05’. The peak summits were then extended by 250 bp on either side to a final width of 501 bp, filtered by the ENCODE hg38 blacklist (https://www.encodeproject.org/ annotations/ENCSR636HFF/), and filtered to remove peaks that extend beyond the ends of chromosomes. We then created a non-overlapping set of extended summits across all of these peaks as described previously^12, 18^.

We then counted the accessibility for each cell in these peak regions to create an accessibility matrix. We then adopted the iterative LSI clustering approach^12, 18^ to unbiasedly identify clusters that are due to biological vs technical variation. Briefly, we computed the TF-IDF transformation as described by Cusanovich et. al.^83^. To do this, we divided each index by the colSums of the matrix to compute the cell “term frequency”. Next, we multiplied these values by log(1 + ncol(matrix)/rowSums(matrix)), which represents the “inverse document frequency”. This yields a TF-IDF matrix that can be used as input to irlba’s SVD implementation in R. We then used Harmony to batch correct the LSI dimensions in R. Using the first 25 reduced dimensions as input into a Seurat object, crude clusters were identified using Seurat’s (v2.3) SNN graph clustering FindClusters function with a resolution of 0.2. We then calculated the cluster sums from the binarized accessibility matrix and then log-normalized using edgeR’s ‘cpm(matrix, log = TRUE, prior.count = 3)’ in R. Next, we identified the top 25,000 varying peaks across all clusters using ‘rowVars’ in R. This was done on the cluster log-normalized matrix rather than the sparse binary matrix because: (1) it reduced biases due to cluster cell sizes, and (2) it attenuated the mean-variability relationship by converting to log space with a scaled prior count. The 25,000 variable peaks were then used to subset the sparse binarized accessibility matrix and recompute the TF-IDF transform. We used SVD on the TF-IDF matrix to generate a lower dimensional representation of the data by retaining the first 25 dimensions. We then used Harmony to batch correct the LSI dimensions in R. We then used these reduced dimensions as input into a Seurat object and crude clusters were identified using Seurat’s (v.2.3) SNN graph clustering FindClusters function with a resolution of 0.6. This process was repeated a third time with a resolution of 1.0. Then, these same reduced dimensions were used as input to Seurat’s ‘RunUMAP’ with default parameters and plotted in ggplot2 using R.

### Identification of clusters and cell types from scATAC-seq data

Different clusters and cell types were manually identified using promoter accessibility and gene activity scores for various lineage-defining genes. Microglia (Cluster 24) were identified based on accessibility near the *IBA1*, *CD14*, *CD11C*, *PTGS1*, and *PTGS2* genes. Astrocytes (Clusters 13-17) were identified based on accessibility near the *GFAP* and *FGFR3* genes. Excitatory neurons (Clusters 1, 3, and 4 were identified based on accessibility near the *SLC17A6* and *SLC17A7* genes. Inhibitory neurons (Cluster 2, 11, and 12) were identified based on accessibility near the *GAD2* and *SLC32A1* genes. Medium spiny neurons (most of Cluster 2) were identified based on accessibility near the *DARPP32* gene. Oligodendrocytes (Clusters 19-23) were identified based on accessibility near the *MAG* and *SOX10* genes. OPCs (Clusters 8-10) were identified based on accessibility near the *PDGFRA* gene. All neuronal subsets, for example nigral neurons (Cluster 5-6), were identified primarily as neurons based on accessibility near the *NEFL*, *RBFOX3*, *VGF*, and *GRIN1* genes and then subdivided based on the region of origin and the accessibility near other genes mentioned above.

### Single-cell ATAC-seq peak calling

For scATAC-seq peak calling from clusters or manually defined cell types, all single cells belonging to the given group were pooled together. These pooled fragment files were converted to the paired-end tagAlign format and processed with version 1.4.2 of the ENCODE DCC ATAC-seq pipeline. The conversion to tagAlign was performed as follows. For fragments on the positive strand, the read start coordinate was the fragment start coordinate, zero-indexed. The read end coordinate was the fragment start coordinate plus the read length (99 bp). For fragments on the negative strand, the read start coordinate was the fragment end coordinate, zero-indexed. The read start coordinate was the fragment end coordinate minus the read length (99 bp). Then, these tagAlign files were used as input to the DCC ATAC-seq pipeline. IDR optimal peak sets with an IDR threshold of 0.05 were determined for each cluster by the pipeline, using pseudo-bulk replicate tagAligns for the cluster. Other pipeline parameters were the same as for bulk ATAC-seq data (see above).

### Single-cell ATAC-seq gene activity scores

We calculated gene activity scores by summing the binarized accessibility, weighted by distance, in the 1-kb tiles within 100 kb. The distance weights were computed by determining the distance from the tile to the gene promoter start site and computing “exp(-abs(distance)/10000)”. These were then scaled to 10,000 and log-normalized with a pseudo count of 1. For visualization purposes, the top and bottom 2.5% of scores were thresholded.

### Single-cell ATAC-seq pseudo-bulk replicate generation and differential accessibility comparisons

For differential comparisons of clusters or cell types, including Pearson correlation determination, non-overlapping pseudo-bulk replicates were generated from groups of cells. For each cell grouping (i.e a cluster or a cell type), a minimum of 300 cells was required in order to make at least two non-overlapping pseudo-bulk replicates of 150 cells each. A maximum of 3 pseudo-bulk replicates was made per group if the total number of cells per group was greater than 450 cells. Cells were randomly deposited into one of the pseudo-bulk replicates and all available cells were used. In this way, the non-overlapping pseudo-bulk replicates are agnostic to which donor the cell came from but aware of individual cells (i.e. all reads from a given cell are deposited into the same pseudo-bulk replicate). These pseudo-bulk replicates were then used for differential comparisons using DESeq2^84^.

### CIBERSORT deconvolution

CIBERSORT^25^ was used to deconvolve bulk ATAC-seq data using signature matrices generated from scATAC-seq data. Default parameters were used. For the cell type-specific classifier, pseudo-bulk replicates were generated for each of the 8 main cell types. For the cluster-specific classifier, pseudo-bulk replicates were generated for each of the 24 clusters.

### Transcription factor footprinting

Transcription factor footprinting was performed as described previously^64^.

### HiChIP library generation

HiChIP library generation was performed as described previously^13^. One million cryopreserved nuclei were used per experiment. Enzyme MboI was used for restriction digest. Sonication was performed on a Covaris E220 instrument using the following settings: duty cycle 5, peak incident power 140, cycles per burst 200, time 4 minutes. All HiChIP was performed using H3K27ac as the target (Abcam ab4729).

### HiChIP data analysis

HiChIP paired-end sequencing data was processed using HiC-Pro^85^ version 2.11.0 with a minimum mapping quality of 10. FitHiChIP^86^ was used to identify “peak-to-all” interactions using peaks called from the one-dimensional HiChIP data. A lower distance threshold of 20 kb and an upper distance threshold of 2 Mb were used. Bias correction was performed using coverage-specific bias.

### HiChIP linkage of SNPs to genes

To link SNPs to genes, we identified FitHiChIP loops that contained a SNP in one anchor and a TSS in the other anchor. This was performed for all LD-expanded SNPs to identify the full complement of genes that could be putatively implicated in AD and PD.

### gkm-SVM machine learning classifier training and testing

For each of the 24 scATAC-seq clusters, we used a 10 fold cross-validation scheme to train weighted gapped k-mer Support Vector Machine (gkm-SVM) models to classify 1000 bp sequences into two classes - accessible (corresponding to sequences underlying peaks) and inaccessible (GC matched inaccessible genomic regions). The test sets for each of the 10 folds are as follows. Fold 0 consisted of chr 1. Fold 1 consisted of chr 2 and chr 19. Fold 2 consisted of chr 3 and chr 20. Fold 3 consisted of chr 6, chr 13, and chr 22. Fold 4 consisted of chr 5, chr 16, and chr Y. Fold 5 consisted of chr 4, chr 15, and chr 21. Fold 6 consisted of chr 7, chr 14, and chr 18. Fold 7 consisted of chr 11, chr 17, and chr X. Fold 8 consisted of chr 9 and chr 12. Fold 9 consisted of chr 8 and chr 10.

For each of the 24 scATAC-seq clusters, we merged the IDR peaks with identical genomic coordinates (peaks with multiple summits) while preserving the summit position and the MACS2 p-value of the peak with the lowest p-value among the ones with the identical coordinates. Next, we ranked the peaks by the MACS2 p-value, expanded each peak by 500 bp on either side of the summit, to a total of 1000 bp, and eliminated those peaks with any ‘N’ bases in the 1000 bp. For each of 10 cross-validation folds, we kept up to 60,000 of the top peaks belonging to the training set and all of the peaks belonging to the much smaller test set, all of which comprised the positively labeled (accessible) examples for training.

In order to generate the negative (inaccessible) examples for each of the cross-validation folds in each single-cell cluster, first, we used seqdataloader (https://github.com/kundajelab/seqdataloader) to generate all 1000 bp sequences obtained by tiling the hg38 genome 200 bp at a time, with a stride of 50 bp, keeping those 200 bp segments that have no IDR peak summits in that cluster, and then expanding those 200 bp segments by 400 bp on each side for a total of 1000 bp. Next, we calculated the GC content of the selected positive examples and all of the negative sequences. We matched each of the positive examples, both in the training set and the test set, with a negative sequence with the closest GC content, without replacement.

For each of the 10 folds in each of the 24 clusters, we used the 1000-bp DNA sequences corresponding to the positive and GC-matched negative training examples as inputs to the gkmtrain function from the LS-GKM package^87^ with the default options, producing a total of 240 models; the default options for LS-GKM included the gapped *k*-mer + center weighted (wgkm) kernel (t = 4), a word length of 11 (l = 11), 7 informative columns (k = 7), 3 maximum mismatches to consider (d = 3), an initial value of the exponential decay function of 50 (M = 50), a half-life parameter of 50 (H = 50), a regularization parameter of 1.0 (c = 1.0), and a precision parameter of 0.001 (e = 0.001). We used the resulting support vectors for each trained model to score the DNA sequences corresponding to the positive and GC-matched negative test set examples for each fold in each cluster by running gkmpredict, and used the scikit-learn python library^88^ to calculate both auROC and auPRC accuracy metrics.

### gkm-SVM allelic scores of candidate SNPs

We intersected the coordinates of all LD-expanded candidate AD and PD GWAS and colocalization SNPs with those of the peaks for each single-cell ATAC-seq cluster to obtain the SNPs in each cluster that are in peaks. For each SNP in a peak in each of the clusters, we retrieved the 1000 bp DNA sequence around the SNP, with the SNP at its center, and created a sequence corresponding to the effect allele by replacing the 500th position of the sequence with the effect allele. Similarly, we created another sequence corresponding to the non-effect allele by replacing the 500th position of the sequence with the non-effect allele. Furthermore, we repeated the same procedure to also produce 50 bp sequences for each SNP with the effect allele and the non-effect allele by retrieving the 50 bp DNA sequence around each SNP and replacing the 25th position with the effect and the non-effect allele, respectively.

For each SNP in a peak in each of the clusters, we computed **GkmExplain**^37^ importance scores for each position in each of the 1000 bp effect and non-effect allele sequences using each of the 10 gkm-SVM^36^ models for the respective cluster. GkmExplain is a method to infer the importance or predictive contribution of every base in an input sequence to its corresponding output prediction from a gkm-SVM model. Next, for each SNP in a given cluster, we computed the average score for each position across all 10 models (from the 10 folds) for that cluster for both the effect allele sequence and the non-effect allele sequence, producing a set of consensus importance scores for both the effect allele and the non-effect allele. Then, we subtracted the sum of these consensus importance scores corresponding to the central 50 bp of the non-effect allele sequence from that of the effect allele sequence to compute the GkmExplain score for each SNP in each cluster.

To compute ***in silico* mutagenesis (ISM)** scores for each SNP in a peak in each of the clusters, we used each of the 10 fold gkm-SVM models from the respective cluster to compute model output prediction scores for the 50 bp effect and non-effect allele sequences by running gkmpredict. Then, we subtracted the score of the non-effect allele sequence from that of the effect allele sequence to obtain the ISM score and computed the average ISM score for each SNP across all 10 folds in each cluster.

To compute **deltaSVM** scores, we generated all possible non-redundant k-mers of size 11 and scored each of them using each of the 240 models. Next, for each SNP in a peak in each of the clusters, we used each of the 10 sets of *k*-mer scores from the gkm-SVM models from the respective cluster to run deltaSVM^39^ on the 50 bp effect and non-effect allele sequences. We computed the average of the resulting deltaSVM scores for each SNP across all 10 folds in each cluster.

### Statistical significance and high confidence sets of gkm-SVM based allelic scores for candidate SNPs

In order to obtain a statistical significance for each of the three gkm-SVM model based allelic SNP scores (GkmExplain, ISM and deltaSVM), we computed an empirical null distribution of scores. We expect most of the LD expanded candidate SNPs to be non functional. Hence, we simply use the distribution of the scores for all candidate SNPs as an empirical null distribution. For each type of score, in order to control for any arbitrary bias in the sign of the score, we included the negative value of each score to the list of scores to enforce symmetry. We found that the t-distribution was a good fit (based on KS test) to the empirical null distribution for all three scores. Hence, we used the fitted t-distributions (using SciPy python library http://www.scipy.org/) to each of the three sets of scores as the null distributions.

To select SNPs with **statistically significant gkm-SVM allelic scores**, for each cluster, we selected those SNPs that fall outside the 95% confidence interval for all three null *t*-distributions fitted to the GkmExplain, ISM, and deltaSVM scores.

Next, we developed a method to identify putative transcription factor binding sites around each gkm-SVM scored statistically significant candidate SNP, by identifying the subsequences around the SNP whose base-resolution importance scores are significantly above background. For each SNP, we defined the **active allele** as the allele for which the 50 bp sequence centered on the SNP has the higher gkmpredict output score (relative to the other allele) from the gkm-SVM model. We fitted a background null t-distribution to the consensus GkmExplain importance scores (averaged across models for all 10 folds) of all bases in the 200 bp sequence centered on the SNP and containing the active allele. We use this null distribution to identify bases around the SNP with high signal-to-noise ratio. Specifically, starting from the center of the positive allele’s sequence, which is the location of the SNP, we continue advancing one pointer upstream and another downstream, each up to the position beyond which lie two consecutive bases that both have consensus importance scores that are within or lower than the 90% confidence interval for the distribution fitted to the consensus importance scores for that sequence. The subsequence between the terminal positions of the two pointers corresponds to one that underlies a series of bases with high GkmExplain importance scores that are significantly above scores of surrounding background seqeunce and potentially contains transcription factor binding sites and motifs that are relevant for the given cluster. We refer to these high-importance subsequences seqlets.

Next, we defined two additional scores (prominence score and magnitude score) to further identify high confidence candidates from the gkm-SVM scored statistically significant candidate SNPs supported by seqlets that could potentially match identifiable transcription factor binding sites. We compute the sum of the non-negative consensus importance scores from the active allele’s seqlet, which we refer to as the **active seqlet score**, and divide that score by the sum of the non-negative consensus importance scores from the entire central 200-bp region of the active allele’s sequence; we refer to this ratio as the **active seqlet signal-to-noise ratio**. Similarly, we compute the **inactive seqlet score** as the sum of the non-negative consensus importance scores in the inactive allele’s sequence from the same positions overlapping the active seqlet. We obtain a corresponding **inactive seqlet signal-to-noise ratio** by dividing the inactive seqlet score by the sum of the non-negative consensus importance scores from the entire central 200-bp region of the inactive allele’s sequence. Then, for each SNP, we compute the **prominence score** by subtracting the non-effect allele’s seqlet signal-to-noise ratio from the effect allele’s seqlet signal-to-noise ratio. In addition, we also compute a **magnitude score** by subtracting the non-effect allele’s seqlet score from the effect allele’s seqlet score.

To compute the statistical significance of the prominence and magnitude scores for candidate SNPs, for each cluster, we fit null *t*-distributions to the prominence scores and magnitude scores (using a KS test to test goodness of fit of the *t*-distribution to the empirical distribution of scores). For each type of score, in order to control for any arbitrary bias in the sign of the score, we include the negative value of each score to the list of scores to enforce symmetry before fitting the distribution.

Finally, to prioritize SNPs that disrupt potential transcription factor binding sites, in each cluster, among the SNPs with statistically significant gkm-SVM allelic scores, we designate as high confidence SNPs those that have prominence scores outside the 95% confidence interval for the distribution fitted to the prominence scores. These are the SNPs that have an allele that completely destroys a prominent and high-scoring seqlet and, as a result, potentially disrupts an important transcription factor binding site. Next, among the confident SNPs that do not pass the high confidence threshold, we designated as medium confidence SNPs those that have either peak magnitude scores outside the 95% confidence interval or prominence scores outside the 80% confidence interval. The magnitude threshold is intended to capture those SNPs that have a significant deleterious effect on the seqlet score, even if those SNPs do not necessarily destroy the entire seqlet and even for cases where the seqlet around the SNP is not among the most prominent seqlets in the local 200 bp sequence window. In addition, the relaxed prominence threshold is intended to capture those SNPs that do not pass the stringent filter for the high confidence set, but nevertheless, demonstrate at least a partial deleterious effect on a moderately scoring seqlet around the SNP. Together, these two filters serve to increase the recall in the prioritization of the SNPs, allowing us to identify all promising SNPs that are worthy of in-depth evaluation, which can assess their potential regulatory effect through a case-by-case analysis. The remaining SNPs in the confident set, which fail to meet the threshold set for medium confidence, are designated as low confidence SNPs, as they include SNPs that significantly reduce the GkmExplain score, the ISM score, and the deltaSVM score, but do not have a clear impact on a seqlet around the SNP, making it unlikely for them to have a disruptive effect on a key transcription factor binding site.

### Identification of MAPT haplotypes

The MAPT haplotype block is part of one of the largest LD blocks in the human genome. To identify SNPs that belong exclusively to either the H1 or H2 haplotype, we used minor allele frequencies from dbSNP version 151. SNPs were required to be within the coordinates of the MAPT inversion breakpoints (hg38 chr17:45551578-46494237) and to have a minor allele frequency between 8.4% and 9%. While there are undoubtedly haplotype specific SNPs outside this frequency range, we chose this range to be as conservative as possible and to pick SNPs that showed minimal haplotype switching. Each SNP was verified to track with the predicted haplotype using LDLink^89^. This resulted in 2366 SNPs that could be confidently called as haplotype divergent.

### MAPT locus differential expression analysis

A 900-kb block of variants in strong LD at the *MAPT* locus hampered the resolution of colocalization methods for identifying causal variants and/or genes at this locus. To probe this locus more deeply, we assembled a list of 2366 variants uniquely found in either the H1 or the H2 haplotype of the *MAPT* locus (described above). For each of the 838 individuals genotyped in GTEx v8, we counted the number of variants in support of either haplotype. We designated individuals as homozygous if they possessed less than 1% of variants favoring the opposite haplotype and heterozygous if 45% to 55% of variants supported either haplotype. This determined the individual’s haplotype in all but six cases, which were excluded from the remainder of the *MAPT* analysis. In total, we identified 539 individuals with the H1/H1 haplotype, 260 with H2/H1, and 33 with H2/H2. Our a priori gene of interest was *MAPT*, which whose expression had previously been demonstrated to be higher in H1 than H2 haplotypes. At a nominal cutoff of p < 0.05, we confirmed this expected direction of differential *MAPT* expression (higher in H1 haplotypes) in multiple tissues, with the strongest contrasts in “Brain - Cortex”.

We then extended our analysis to include all genes expressed in any of the brain tissues from GTEx v8. We compared the log2-fold change of gene expression (TPM) between H1/H1 and H1/H2 individuals, given that these subgroups had the largest sample size. A change was considered statistically significant if a Wilcoxon rank-sum test between the two groups produced a p-value of < 0.05 / (total # genes) / (total # tissues). We also performed pairwise Wilcoxon rank-sum test comparisons for each gene in each brain tissue between all 3 pairings of haplotypes.

### MAPT haplotype-specific ATAC-seq and HiChIP analysis

For both ATAC-seq and HiChIP, reads from heterozygote donors were re-mapped to an N-masked genome (using bowtie2 or HiCPro, respectively) where all dbSNP v151 positions were masked to “N”. After alignment, SNPsplit^90^ was used to divide reads mapping to either the H1 or H2 haplotypes based on the presence of one of the 2366 haplotype-divergent SNPs identified above. In this way, reads mapping to regions that lack a haplotype-divergent SNP could not be assigned in an allelic fashion to either the H1 or H2 haplotypes and were ignored. For track-based visualizations of haplotype-specific data, all available data from a given haplotype was merged agnostic to what brain region the data was derived from. To identify regions with haplotype-specific chromatin accessibility in the MAPT locus, the entire locus was tiled into non-overlapping 500 bp bins and the number of Tn5 transposase insertions were counted for each haplotype in each bin for each sample. A Wilcoxon signed-rank test was used to determine if the difference between H1 and H2 for each bin was significant after multiple hypothesis correction (FDR < 0.01).

